# Mitochondrial mutation spectrum in Chordates: damage versus replication signatures, causes, and dynamics

**DOI:** 10.1101/2023.12.08.570826

**Authors:** Dmitrii Iliushchenko, Bogdan Efimenko, Alina G. Mikhailova, Victor Shamanskiy, Murat K. Saparbaev, Ilya Mazunin, Dmitrii Knorre, Wolfram S. Kunz, Philipp Kapranov, Stepan Denisov, Jacques Fellay, Konstantin Khrapko, Konstantin Gunbin, Konstantin Popadin

## Abstract

To elucidate the primary factors shaping mitochondrial DNA (mtDNA) mutagenesis, we derived a comprehensive 192-component mtDNA mutational spectrum using 86,149 polymorphic synonymous mutations reconstructed from the CytB gene of 967 chordate species. The mtDNA spectrum analysis provided numerous findings on repair and mutation processes, breaking it down into three main signatures: (i) symmetrical, evenly distributed across both strands, mutations, induced by gamma DNA polymerase (about 50% of all mutations); (ii) asymmetrical, heavy-strand-specific, C>T mutations (about 30%); and (iii) asymmetrical, heavy-strand-specific A>G mutations, influenced by metabolic and age-specific factors (about 20%). We propose that both asymmetrical signatures are driven by single-strand specific damage coupled with inefficient base excision repair on the lagging (heavy) strand of mtDNA. Understanding the detailed mechanisms of this damage is crucial for developing strategies to reduce somatic mtDNA mutational load, which is vital for combating age-related diseases.

## Introduction

DNA mutations can be a result of either replication error or damage followed by incorrect repair^1^, with a range of mutagens responsible for these changes^2^. Reconstruction of a mutational spectrum, with 96 or 192 components, helps to decompose it into distinct mutational signatures, allowing us to trace the effects of various mutagens^3^. In the human nuclear genome, both germline and somatic spectra, including cancerous ones, have been instrumental in reconstructing and deconvoluting these signatures, leading to major breakthroughs in both fundamental^4^ and applied^5^ research.

Despite significant advancements in comprehending the mutagenesis of the human nuclear genome, the mitochondrial genome is less well-characterised, yet playing a vital role in numerous human diseases^6^ and ageing^7^. The mutagenesis of the mitochondrial genome is mysterious: being a hundred times faster than in the nuclear genome, it remains partially elucidated, since its spectra show neither expected mutational signatures of reactive oxygen species (ROS), UV light nor tobacco smoke in associated cancer data^8^. Furthermore, the mechanisms of mtDNA replication and repair have been under ongoing debates, and recent detailed analysis of the mitochondrial mutational spectrum offered important insights into these processes^9^. On an evolutionary scale, understanding mtDNA mutagenesis and implementation of mtDNA spectrum-aware approaches is crucial for accurate phylogenetic inferences and for uncovering variations in selection processes between species^3,10^ as well as for understanding the dynamics of deleterious human mtDNA variants.

The reconstruction of a detailed mutational spectrum of single base substitutions for each species demands extensive data. Traditionally, comparative species analyses employ the transitions to transversions ratio as a basic characteristic of the mutational spectrum^11^. Availability of more sequence data, allowed researchers to use more complex 6-component (focusing only on pyrimidines in Watson-Crick base pairs: C>A, C>G, C>T, T>A, T>C, T>G, under the assumption of symmetrical mutagenesis on complementary strands)^12–14^ and 12-component (doubling the 6-component spectrum to account for asymmetrical mutagenesis on complementary strands)^10^ spectra in analyses for comparative studies. However, more comprehensive 96-component (expanding the 6-component spectrum with an inclusion of the nucleotide context) and 192-component (a doubled version of the 96-component spectrum, assuming asymmetry between complementary strands) spectra, which consider adjacent nucleotide context, have been limited to extensively studied species such as humans^8^ and SARS-CoV-2^15^. For rarely-sequenced, non-model taxa, constructing these in-depth spectra is challenging.

Here, focusing on chordates, we overcame this limitation by integrating rare, species-specific mtDNA polymorphisms into a comprehensive spectrum representative of all *Chordata*. Reconstructing 86,149 synonymous polymorphic mutations from CytB sequences of 967 chordate species, we compiled a 192-component mtDNA spectrum. This extensive spectrum enables the exploration of critical questions, related to damage and replication-driven mechanisms of mtDNA mutagenesis, their etiologies, and dynamics.

## METHODS

### In this paper we use heavy strand notation for all mtDNA mutations and spectra

#### 1. Reconstruction of the integral 192-component mutational spectrum for CytB for all chordates

To reconstruct the mutational spectrum for CytB, we made the assumption that the majority of mtDNA synonymous polymorphic variants within chordate species are effectively neutral. Although recent suggestions indicate that certain synonymous sites in mtDNA may not be entirely neutral^16^, we evaluated the effect of removing highly-constrained synonymous sites on our data. The results suggested that omitting these conservative sites did not markedly change the results (see Supplementary Figure 1).

We followed these steps:

1a) Observed mutations

Only species with at least five unique sequences of CytB gene from GenBank were used in this work. For each of 967 such species (listed in Supplementary Table 1) the observed mutational spectrum was derived through the following steps: (i) obtaining a codon-based multiple alignment using macse v2^17^; (ii) reconstructing the phylogenetic tree and rooting it with the closest relative species as the outgroup; (iii) reconstructing ancestral sequences (most probable nucleotide in each alignment position) at each internal node using RAxML^18^; (iv) identifying all single-nucleotide synonymous substitutions on each tree branch; (v) categorising these mutations into 192 groups based on the nucleotide context (16 unique contexts of downstream and upstream nucleotides within each of 12 mutation types).

1b) Expected mutations

To consider differences in nucleotide and trinucleotide composition, for each of 967 species we calculated the expected mutational spectra. Focusing on the reference CytB gene, obtained from GenBank for each analysed chordates species, we executed an *in silico* saturation mutagenesis procedure. The central concept behind *in silico* saturation mutagenesis is to replace each nucleotide with one of three possible alternatives and select only synonymous substitutions. These substitutions were then recorded alongside their neighbouring nucleotides, and categorised according to the specific type of substitution they represented.

1c) To obtain species-specific mutational spectra

We normalised observed mutations to expected ones. For each species, we calculated the mutation rates by dividing the number of observed substitutions by the number of expected substitutions in the corresponding context, resulting in 192 substitution rates. These rates were then transformed into frequencies, ensuring that the sum of all 192 normalised substitution rates equaled 1 for each species.

1d) Integral taxa-specific mutational spectrum

Finally, we averaged the mutational spectra of species samples to derive class-specific or phylum-specific mutational spectra.

#### 2) Comparison of the mutational spectra

We primarily used cosine similarity to estimate the closeness of the compared spectra. Comparison has been performed on two levels: between species and between classes. Firstly, we conducted pairwise comparisons of mutational spectra at the species-specific level within and between classes, considering all possible combinations. Second, we estimated differences between classes using jackknife resampling of species spectra. In this process, we randomly selected 20 species from each pair of classes, calculated the 192-component mutational spectrum for both classes, and computed the cosine similarity of either the overall mutational spectrum or its parts (transitions and transversions). We repeated this process in 1000 iterations for every conceivable class combination. Through the use of jackknife resampling, we effectively adjusted the influence of varying class sizes.

#### 3) Annotation of mitochondrial mutational signatures

To decompose class-specific mtDNA mutational spectra into COSMIC signatures we used SigProfilerAssignment tool v0.0.30^19^. Since main COSMIC signatures are symmetrical and do not account for strand-specific mutagenesis, we split the asymmetrical mitochondrial 192-component spectra into two (“low” and “high”) 96-component spectra based on the abundance of specific transitions. The “high” spectra include more frequent C_H_>T_H_ and A_H_>G_H_ transitions, while the “low” spectra include G_H_>A_H_ and T_H_>C_H_ transitions, where H signifies heavy strand notation of substitution. Also, to analyse the asymmetrical component of mutagenesis, we derived “diff” spectra subtracting “low” from “high” spectra in a context-dependent manner (Supplementary Fig. 6). All complementary pairs of transversions rates (for example, A_H_>C_H_ and T_H_>G_H_) were averaged and equally added to the “low”, “diff” and “high” spectra; otherwise, to test the impact of noisy and rare transversions, they were zeroed in these spectra. In addition, SigProfilerAssignment, like any other SigProfiler tool, is capable of processing distinct substitution counts on the reference human nuclear genome. To ensure correspondence of input spectra to the human nuclear genome, we rescaled spectra of analysed classes multiplying them by the trinucleotide frequencies of the human nuclear genome. We applied SigProfilerAssignment using the cosmic_fit function, with the following parameters: genome_build=’GRCh37’, nnls_add_penalty=0.01 (reduce number of derived noisy signatures that explain low number of mutations), cosmic_version=3.3. Furthermore, to reduce the effect of the unexpected signatures in decomposition of average mammals spectra, we excluded the following subgroups of signatures from the analysis: immunosuppressants, treatment, colibactin, lymphoid, and artefact.

#### 4) Abasic sites patterns in mtDNA

The distribution of abasic sites (AP) at single-nucleotide and single strand (separately for light and heavy) resolution of in mouse mitochondrial genome (mm10) was obtained from Cai et al.^20^ We quantified the AP sites occurrences within all 64 trinucleotide sequences of each strand of mouse mtDNA, excluding the control region. These values were then adjusted by the counts of trinucleotide motifs in a strand-specific manner.

We created trinucleotide logos using the Python library logomaker^21^. These logos used the normalised AP sites count for each trinucleotide as the nucleotide weights. These weights were averaged across all 64 trinucleotides, and final nucleotide counts at each trinucleotide position were normalised and converted to frequencies.

#### 5) Calculation of the mtDNA mutational spectrum asymmetry

The assessment of mtDNA asymmetry involved the transformation of a 192-component mutational spectrum into a 96-component spectrum. This was achieved by selecting frequencies of 96 single base substitutions of pyrimidines only (C>A, C>G, C>T, T>A, T>C and T>G) from the mitochondrial DNA mutational spectrum (see COSMIC) and dividing them by complementary substitutions frequencies.

The total mitochondrial asymmetry was determined by summing the differences between mutations presented in the 96-component mitochondrial mutational spectrum and their complementary mutations.

#### 6) Analysis of mtDNA mutational spectrum in human cancers

Mutations in the full human mtDNA were derived from comprehensive analysis of the human mitochondrial genome by Yuan et al.^8^ The mutational spectrum was calculated as described above, using the CRS reference sequence NC_012920.1. We assumed that almost all mtDNA mutations in cancer are nearly neutral and employed all mutations, including non-synonymous ones, to calculate the nearly neutral spectrum.

In our analyses, we calculated spectra for different parts of mtDNA. To compare the mutational spectra of genome regions that have low and high time spent single-stranded (TSSS) due to asynchronous replication, we separately calculated spectra for the region with low TSSS (first half of the major arc, 5,800 - 10,800) and the region with high TSSS (second half of the major arc, 11,000 - 16,000).

#### 7) Data and code availability

All analyses we performed in python and R. Scripts and data are available on GitHub: https://github.com/mitoclub/mtdna-192component-mutspec-chordata.

## RESULTS

### Variations in mtDNA Mutational Spectrum within Chordates: Importance of Damage

The 192-component mitochondrial mutational spectrum (Fig. 1, Supplementary Table 1) was generated by integrating 86,149 synonymous mutations obtained from CytB sequences across 967 chordate species. Following the exclusion of highly-constrained synonymous sites which did not significantly affect the overall results (Methods, Supplementary Fig. 1) we assume that our mutational spectrum can be considered neutral. The resulting spectrum highlights a significant prevalence of transitions, with C_H_>T_H_ and A_H_>G_H_ being the most frequent types (Fig. 2a). This mutational pattern mirrors findings observed in mammalian germline mutations^10^, somatic mutations in human cancers^8^, and healthy tissues^22^ (Supplementary Table 2).

**Figure 1.**
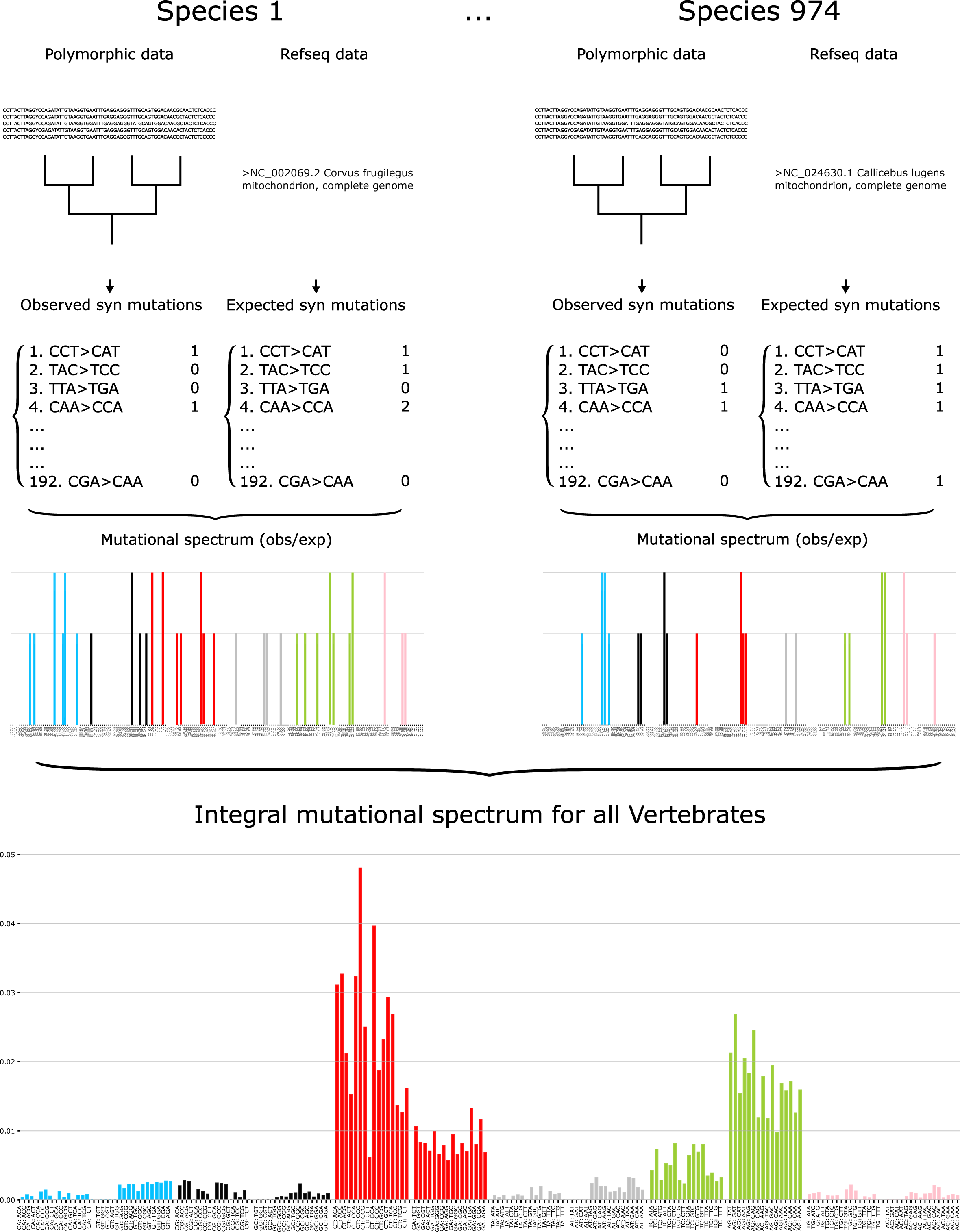
Pipeline overview. For 967 chordate species two groups of mutations were obtained from CytB gene: observed synonymous mutations (from our polymorphic database with total number of substitutions 86,149; Species 1, left column) and expected synonymous substitutions (based on NCBI RefSeq database; Species 1, right column). Both groups of mutations were normalised and integrated into a comprehensive 192-component spectrum for all chordates (Methods).

**Figure 2.**
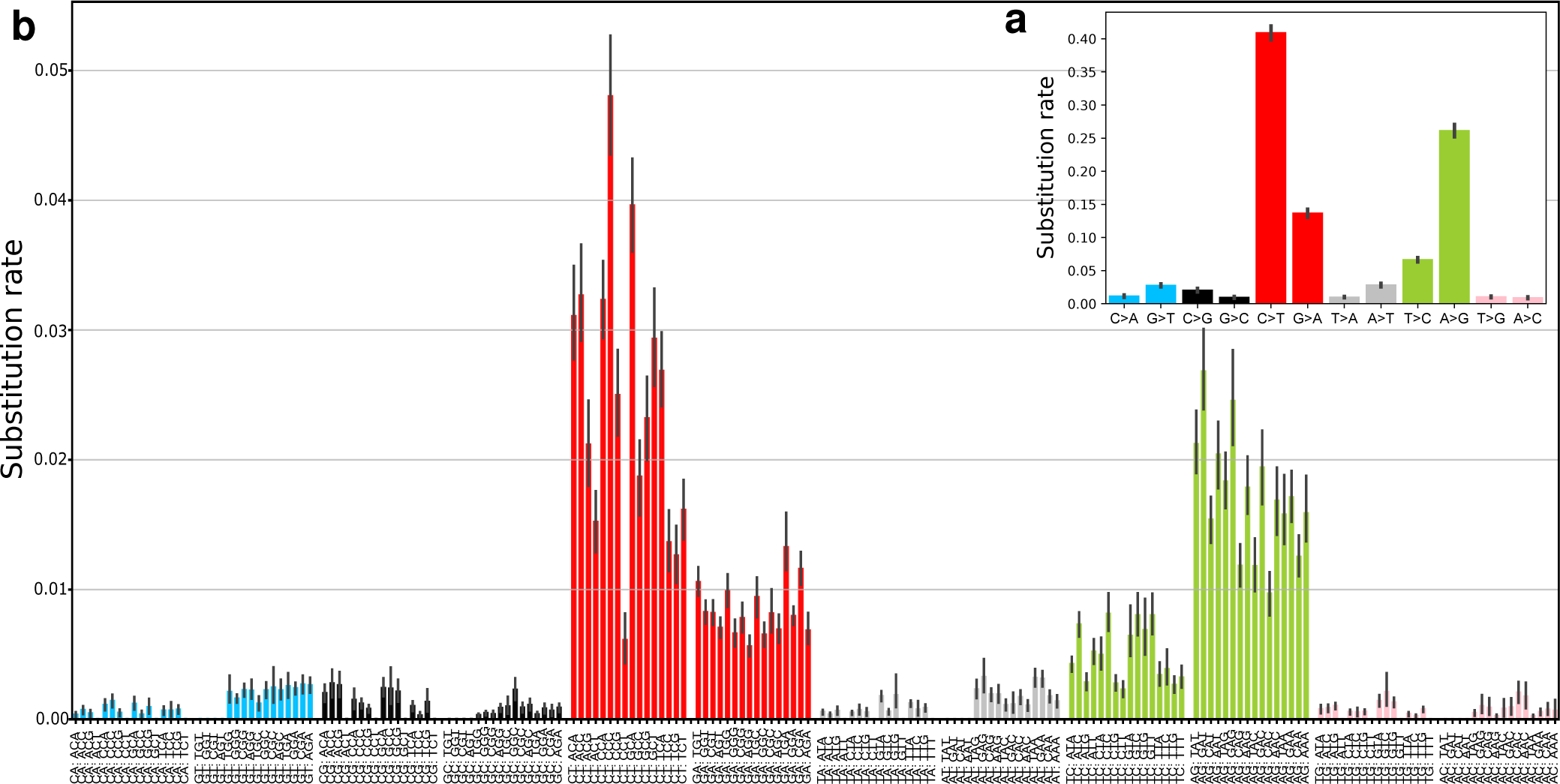
Integral mtDNA mutational spectrum of Chordata. (a) 12-component mutational spectrum (n=967); (b) 192-component mutational spectrum (n=967). The order of substitutions is based on reverse complemented mutations, where, for example, the third bin for C>A substitutions is represented as ACG, and the third bin for G>T substitutions is CGT. Missing (zero) bins are explained by the absence of observed synonymous substitutions, the absence of expected substitutions or both. (c) A scheme, visualising symmetrical and asymmetrical parts of a given mutation.

To assess variation in spectra, we calculated pairwise cosine similarities between all species and revealed no clustering of species spectra within classes (Supplementary Fig. 2). The lack of similarities within class-specific spectra points to fast evolutionary changes in the mtDNA mutation spectra^2^. Such changes can be driven either by variable replication-driven component (if POLG’s properties vary strongly between species) or variable damage-driven component (if mtDNA damage is affected by the metabolism-related traits which can be highly variable even among close species). Taking into account the high evolutionary conservatism of chordate POLG^23–25^ and rather variable levels of aerobic metabolism in different chordates^26^ we propose that the damage effect can be the main reason for the observed high variation. To test it we performed class-specific analyses, expecting that aves, a class characterised on average by the highest level of basal metabolic rate (BMR)^26^, can show the most divergent patterns of the spectrum. We reconstructed mutational spectra for each of the five chordate classes (Supplementary Fig. 3), revealing notable similarities across them and to the overall mtDNA mutational pattern (Fig. 2b), suggesting a conserved mutational process within chordates. Further analysis, involving median pairwise cosine similarities between different classes, confirmed that birds (aves) possess the most distinct mtDNA mutational spectrum among all classes (Methods; Fig. 3a), with the lowest cosine similarities to all other chordate classes. This is in line with a hypothesis that the mtDNA mutational spectrum is shaped by metabolism-associated chemical damage^27^.

**Figure 3.**
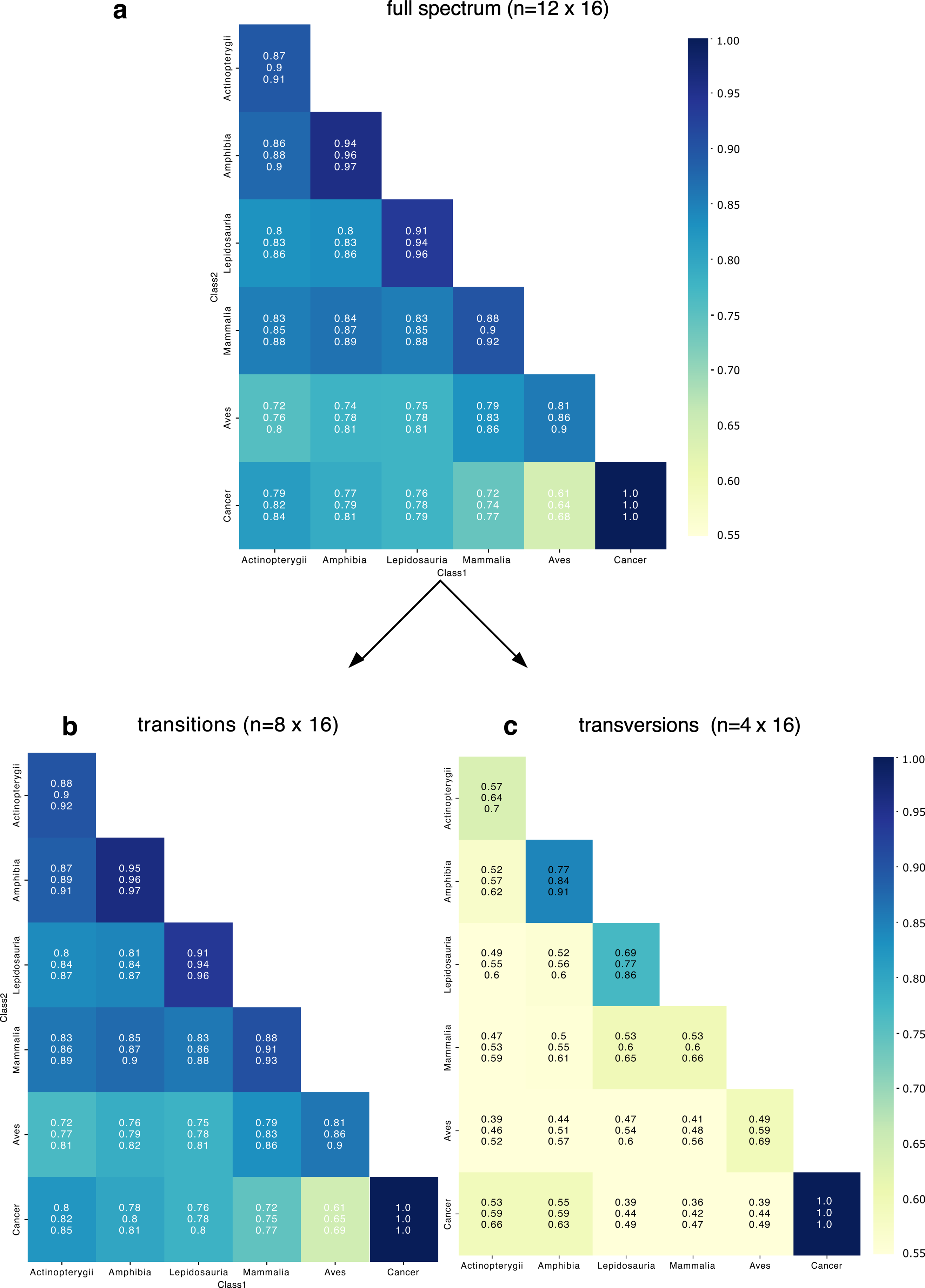
Cosine similarities in pairwise comparisons of somatic and germline variants across five chordate classes and human cancer. Each box presents three values: Q1, Q2 (median), and Q3, which were derived from 1000 cosine similarity comparisons between two classes. (a) High median cosine similarities (0.65 or higher) were consistently observed across five classes of chordates and human cancer when comparing the whole 192-component mutational spectrum (n=192). (b) Similarly, high median cosine similarities (0.83 or higher) were observed specifically for transitions alone (n = 4×16). (c) In contrast, low median cosine similarities (less than 0.6) were observed exclusively for transversions (n = 8×16).

To further test the potential damage effect, we analysed somatic mtDNA mutations from human cancers, which are often hypoxic^28^ and, thus, may have less oxidative damage^29^. Conducting pairwise comparisons between human cancers^8^ (Methods) and five chordate classes we indeed revealed the least cosine similarity between birds and cancers (Fig. 3a), indicating their strong deviations from each other presumably due to the lowest damage in cancers and the highest damage in birds^30,31^. Altogether we propose that due to the differences in the level of aerobic metabolism all chordate species and classes as well as human cancers are affected by mtDNA damage of differential impact.

### Conservative and variable patterns of the mtDNA spectrum: symmetrical replication-driven and asymmetrical damage-driven mutations

To differentiate between stable and variable mtDNA mutational patterns across classes, we analysed transitions and transversions separately. Transitions showed consistently high similarities among chordate classes and human cancers, aligning with the full spectrum trends (Fig. 3b). Conversely, transversions presented lower similarities, likely due to their stochastic occurrence and rarity in our dataset (Fig. 3c; Supplementary Fig. 4). Subsequently, we focused on an analysis of each transition type separately.

Cosine similarities across classes were uniformly high for each of the four transition types (Fig. 4), yet interesting variations were observed. Notably, G_H_>A_H_ exhibited higher similarity between classes than C_H_>T_H_, despite being complementary equivalents. This indicates that symmetrical C>T mutations on double-stranded DNA (dsDNA) (approximated by G_H_>A_H_, Fig. 2c) are more conserved compared to the asymmetrical part of C>T mutation in single-stranded DNA (ssDNA) (approximated by the difference between C_H_>T_H_ and G_H_>A_H_, Fig. 2c). We suggest that the conserved pattern of symmetrical C>T substitutions (equal to G_H_>A_H_) primarily results from internal replication errors, introduced by POLG. Conversely, the less conserved pattern of asymmetrical C>T mutations (C_H_>T_H_ minus G_H_>A_H_) on ssDNA, particularly divergent between aves and cancer, may be influenced by chemical damage including spontaneous deamination or oxydation.

**Figure 4.**
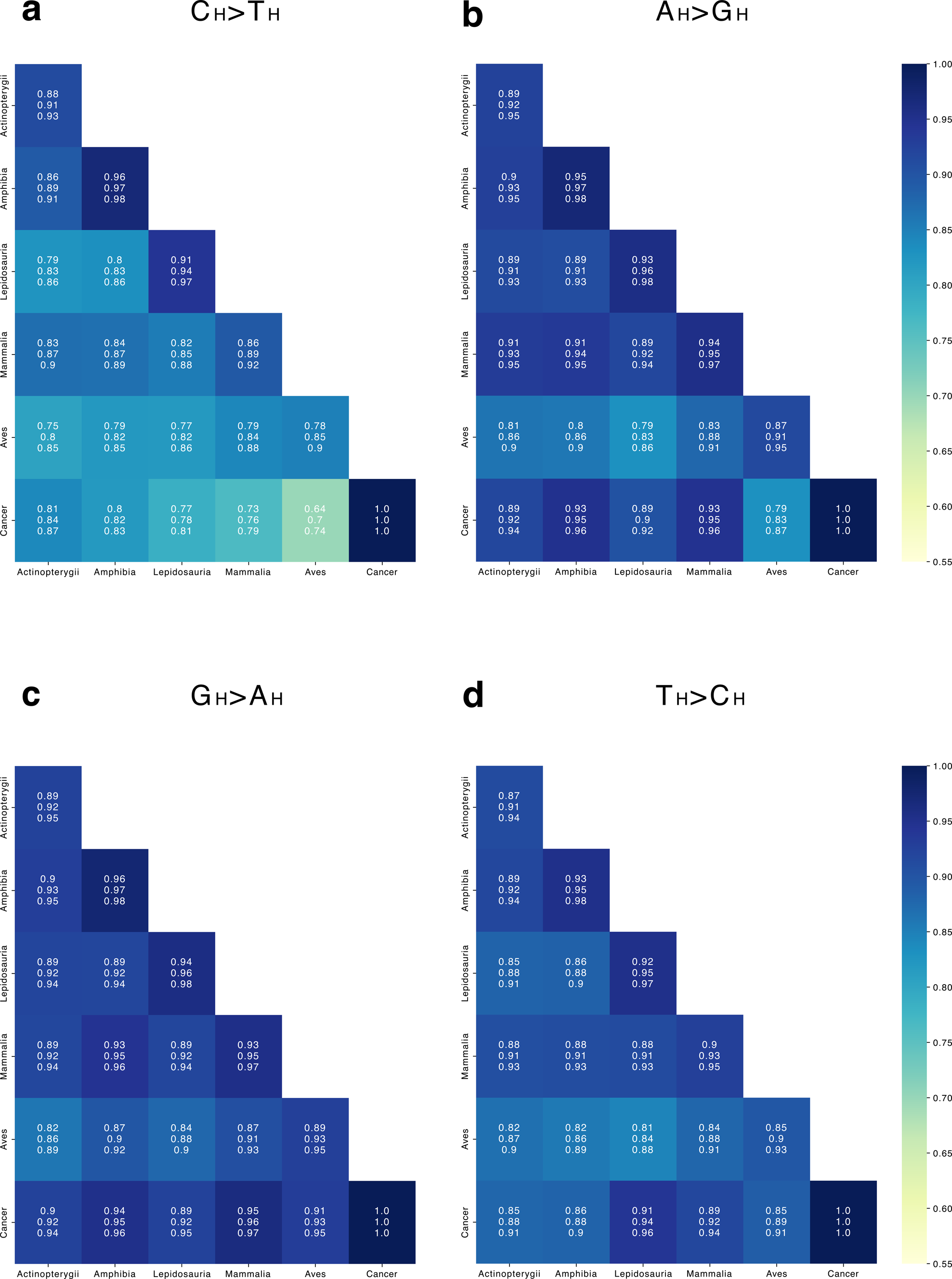
Comparative cosine similarity analysis of transition types across chordate classes and human cancer. Building on the conceptual framework of Figure 3, this figure delineates the cosine similarities for four transition types - (a) C_H_>T_H_, (b) A_H_>G_H_, (c) G_H_>A_H_, and (d) T_H_>C_H_ - across five chordate classes and human cancer mutations. Each transition type is analysed with all possible nucleotide contexts (n=16). The results indicate a high degree of similarity for all transition types examined. Notably, the mutational base G_H_>A_H_ exhibits more conservation compared to the C_H_>T_H_ substitution.

Taking into account that asymmetrical C_H_>T_H_ (i.e. C_H_>T_H_ minus G_H_>A_H_) is most likely associated with chemical damage such as spontaneous deamination or oxidation at ssDNA^9^, we explored the profile of the C_H_>T_H_ substitutions in detail. A yeast experiment with ssDNA and ROS (oxygen peroxide and paraquat) demonstrated that cCc>cTc mutations are a primary target of oxidative damage in ssDNA^32^. Notably, completely in line with these experimental results, we consistently identified cCc>cTc as the most common motif for C_H_>T_H_ substitutions in all chordate classes (Supplementary Fig. 5a). Therefore, the cCc>cTc mutation, the most prevalent in mtDNA across all chordates, likely arises not only from spontaneous deamination but also includes contributions from oxidative or other types of damage to ssDNA. Conversely, cancer data again demonstrates a distinct pattern: asymmetrical C_H_>T_H_ substitutions predominantly appear in the nCg context rather than cCc (Supplementary Fig. 5c), suggesting influences from potentially lower levels of oxidative damage^7^ or from cytosine methylation in mtDNA^33^, paralleling the well-known CpG > TpG substitutions in the nuclear genome.

Analysing the local patterns of A>G substitutions we observed that nAt>nGt and nAg>nGg are the most common motifs across all chordates (Supplementary Fig. 5b). Conversely, it alters in cancers, demonstrating once again diverse environments in somatic cancer cells relative to germ-line tissues (Supplementary Fig. 5d). The A_H_>G_H_ mutation, recently linked to ageing in mammals^10^ and body temperature in chordates^34^, suggests a signature of damage related to aerobic metabolism^10^, yet the exact process driving these substitutions is still unknown.

The potential link to this mutation can be due to the N6-methyldeoxyadenosine (6mA) in mtDNA, being strongly enriched on the heavy strand^35^ and associated with stressful hypoxic conditions. Although enrichment in 6mA in mammalian mitochondria is still contradictory^36,37^ and a link between 6mA and A_H_>G_H_ is rather suggestive, we observed that the 6mA motifs described as (c/a)At and A(t/g) in previous studies^38,35^ are similar to our A_H_>G_H_ motifs nAt>nGt and nAg>nGg (Supplementary Fig. 5b). Future studies are needed to uncover a mechanism behind A_H_>G_H_ substitutions in mtDNA.

Altogether, splitting mtDNA spectrum into conservative and variable parts we propose that the symmetrical part (C_H_>T_H_ equal to G_H_>A_H_) can be shaped by replication-driven POLG-induced mutations while the asymmetrical part (C_H_>T_H_ minus G_H_>A_H_ and A_H_>G_H_ minus T_H_>C_H_) is most likely shaped by damage-driven mutations, induced by deamination, oxidation, or methylation.

### mtDNA mutations through the lens of COSMIC signatures: BER deficiency and *C_H_>T_H_*, MMR absence and G*_L_*>A*_L_*, and SBS12-driven A*_H_*>G*_H_* alterations

To deconvolute the overall mutational spectrum of mtDNA into its underlying mutational signatures, we utilised the COSMIC SBS database (https://cancer.sanger.ac.uk/signatures/sbs/). Since the COSMIC SBS database is built upon 96 component signatures, we divided our 192-component mtDNA spectrum into three sets (Supplementary Fig. 6; Methods): “high” spectrum for both symmetric and asymmetric transitions (substitutions with the highest rate out of two complementary substitutions: nCn>nTn and nAn>nGn), “low” spectrum for symmetric transitions (substitutions with the lowest rate out of two complementary substitutions: nGn>nAn and nTn>nCn), and “diff” spectrum, typical for the heavy strand ssDNA, calculated by subtraction of “low” spectrum from “high” (nCn>nTn minus nGn>nAn and nAn>nGn minus nTn>nCn). All transversions were complementary averaged and added to each set. Given the rarity and variability of transversions in mtDNA, we conducted analyses with all substitutions and separately with only transitions, which confirmed that excluding transversions does not significantly change our findings. We observed that five signatures, namely SBS30, SBS44, SBS21, SBS5 and SBS12 are predominant in mtDNA mutations with varying contributions to high, low and diff spectra (Fig. 5a). SBS5 shows a consistently uniform signature, more pronounced when including transversions.

**Figure 5.**
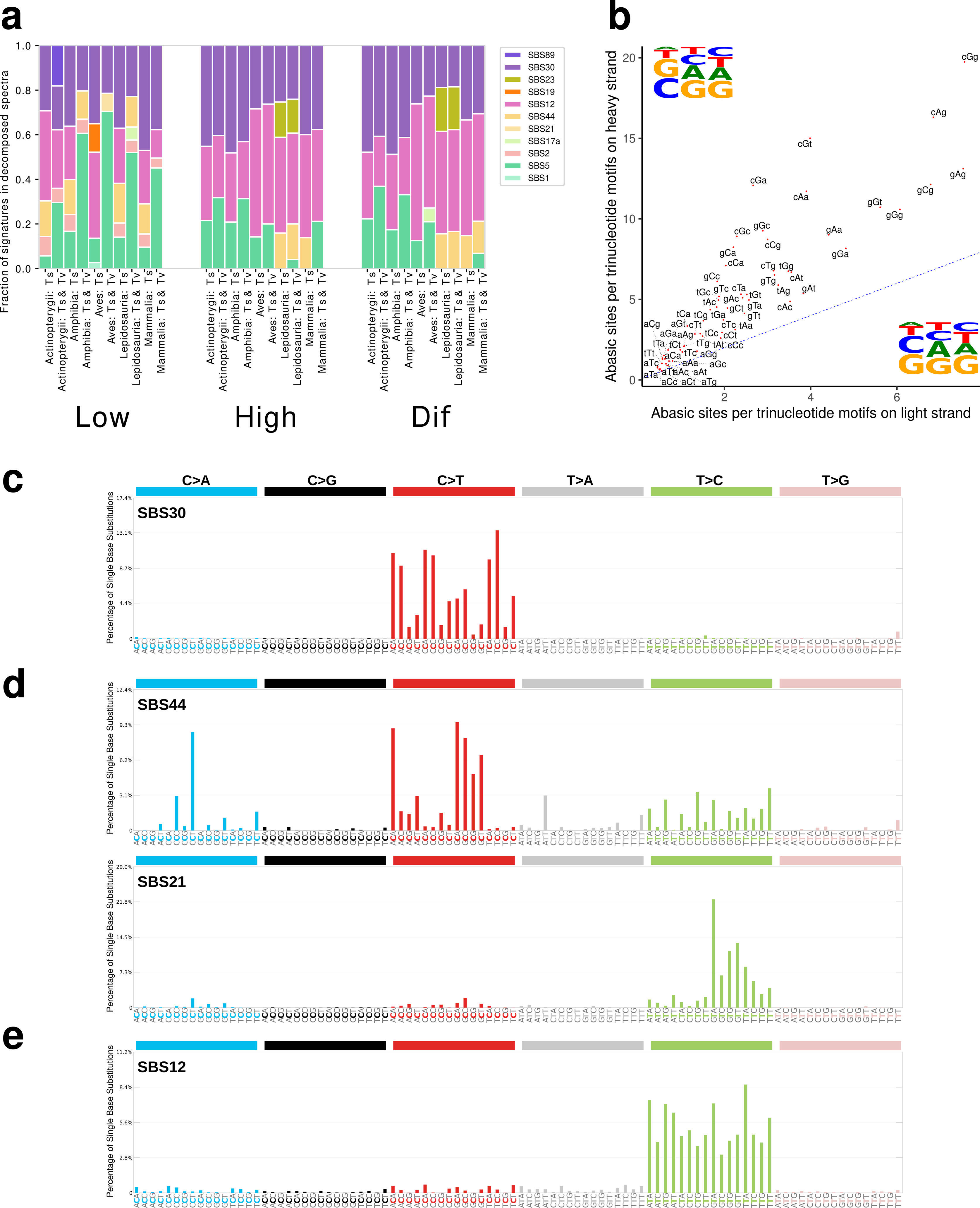
The 192-component mutational spectra deconvoluted into COSMIC SBS signatures. (a) Five signatures SBS30, SBS44, SBS21, SBS5, and SBS12 predominate in spectra within three distinct mutation sets: high, low and diff, as detailed in the main text and Methods section. (b) Analysis of the trinucleotide pattern of AP sites within coding sequences of both the heavy and light strands reveals that trinucleotides on the heavy strand exhibit higher levels of damage, with G being the most frequently damaged nucleotide. (c) Pattern of SBS30 signature: BER deficiency mutations^63^. (d) Pattern of SBS44 and SBS21 signatures: MMR deficiency mutations^63^. (e) Pattern of SBS12 signature: ssDNA-specific mutations^63^.

SBS30 is associated with deficient base excision repair (BER) in the nuclear genome and predominantly results in C>T mutations^39,40^. In mitochondria, BER is the primary repair pathway for chemically damaged bases, including deaminated, oxidised, and alkylated bases^41^. Our findings indicate that in comparison with the nuclear genome, BER in mtDNA is less efficient, and leads to C_H_>T_H_ mutations, particularly in conditions of ssDNA (”high” and “diff” spectra, see Fig. 5a), when there is no complementary strand to replace removed damaged bases. Another notable observation is the association of SBS30 in the nuclear genome with malfunctioning NTHL1 glycosylase, suggesting an inherent decreased efficiency of this enzyme in mtDNA.

NTHL1 removes oxidised pyrimidine lesions in both nuclear DNA (nDNA)^42^ and mtDNA^43^; significantly decreased NTHL1 activity in Friedreich’s ataxia leads to the accumulation of mtDNA mutations dramatically affecting mitochondrial functioning^44^. These observations suggest that NTHL1 could serve as a strategic target for mitigating the mitochondrial mutation rate in clinical settings.

Further insights into mtDNA mutagenesis can be obtained by examining abasic sites (AP sites), which emerge either as an intermediate step of the BER process or spontaneous base loss. In cases where BER is ineffective on ssDNA and there is no glycosylase activity, AP site motifs resembling SBS30 patterns (C_H_>T_H_ mutations) are not expected. Spontaneous base loss will show distinct tendency: depurination (A and G loss) outpaces depyrimidination (C and T loss) by 20-fold, guanines are 1.5 times more prone to depurination than adenines, depurination happens faster in ssDNA than in dsDNA^45^. Analysing AP sites in mouse mtDNA, with precise single-nucleotide resolution on both strands^20^, we observed that motifs with AP sites correlate well between the light and heavy strands, occurring about twice as often on the heavy strand; with Guanine being the most common AP site, followed by Adenine (Fig. 5b). This pattern is more consistent with spontaneous base loss, not BER. Consequently, due to BER deficiency, we do not detect AP sites corresponding to SBS30 motifs. Rather, we find motifs indicative of spontaneous base loss (G and A loss). This process, upon replication by POLG that incorporates a dA residue opposite the abasic site^46^, can result in G>T and A>T transversions. The asymmetry observed for G>T and A>T transversions (Fig. 2a) goes in line with a predominance of AP sites on the heavy strand (Fig. 5b).

SBS44 and SBS21 are two of seven known signatures linked to defective DNA mismatch repair (MMR), crucial for correcting mismatches during DNA replication^47^. Although the presence of MMR in mtDNA has been debated, it is now widely accepted that there is no MMR in mtDNA^41^. Our analysis confirms it showing, that even if MMR partially exists in mtDNA, it is highly deficient. Notably, a pronounced MMR deficiency signature appears in the “low” spectra, associated with symmetrical mutations consistent with symmetric polymerase errors (Fig. 5a). We suggest that MMR-deficiency signatures, i.e. the symmetrical part of C>T (equal to G_H_>A_H_) (Fig. 5d) is shaped by the gamma DNA polymerase, which is expected to introduce symmetrical mutations^40,46,48^.

SBS12, despite its unknown origins in the nuclear genome, can be a hallmark of chemically damaged adenines^22^ in mtDNA’s single-stranded heavy strand due to several reasons. First, SBS12 is based on A_H_>G_H_ mutations (T>C in COSMIC notation, Fig. 5e), which in mtDNA are sensitive to age and temperature^10,49^. Second, SBS12 is presented in the “high” and “diff” spectra (Fig. 5a), reflecting its impact on the heavy strand. Third, SBS12 shows: an increase with replication timing in the nuclear genome (“Replication timing” section in COSMIC), transcriptional strand asymmetry with more A>G mutations on the non-transcribed strand (more T>C mutations on transcribed strand in COSMIC notation) and replication strand asymmetry with A>G mutations on the lagging strand (T>C on leading strand in COSMIC notation). Fourth, SBS12 has the highest prevalence in birds (Fig. 5a), known for elevated metabolic rates, which further underscores its association with chemical damage. SBS12 therefore offers great promise for uncovering the mechanism of mitochondrial ssDNA-specific damage.

Our analysis highlights three primary mutational signatures in mtDNA: (i) Symmetrical C_H_>T_H_ (and complementary G_H_>A_H_) mutations, reflecting mtDNA polymerase errors due to the lack of MMR in mtDNA; (ii) Asymmetric C_H_>T_H_ mutations, likely resulting from cytosine modifications coupled with inefficient BER on ssDNA; (iii) Asymmetric A_H_>G_H_ mutations, linked to metabolism-related adenine damage on ssDNA.

### Strong asymmetry in mtDNA mutagenesis is shaped by single-strand DNA damage coupled with Base Excision Repair deficiency during asynchronous replication

Throughout our paper we have repeatedly demonstrated the pronounced asymmetry of mtDNA mutagenesis: (i) the most common transitions C>T and A>G occur several times more often on heavy strand; (ii) transversions G>T, A>T and C>G also demonstrate increased frequencies on heavy strand in our (Fig. 2a) and other studies^8^; (iii) both mostly pronounced signatures, BER deficient SBS30 and mito-specific SBS12, are highly asymmetrical with a much stronger impact on a heavy strand (Fig. 5a-b). Estimation of the total level of asymmetry in mtDNA (Methods) shows that approximately 50% of all mtDNA mutations are asymmetrical, i.e. occur exclusively on a heavy stand, while the rest of mutations occur symmetrically on both heavy and light strands. Here, we analyse deeper the phenomenon of asymmetry to identify its primary causes.

In the nuclear genome, two types of mutational asymmetry: T-asymmetry (transcription asymmetry, which originates from mutations on the non-transcribed strand) and R-asymmetry (replication asymmetry, which predominantly occurs on the lagging strand) have been described and a tight correlation between both of them has been shown^4^. We calculated mito-asymmetry from our 192-component spectra (Methods) and observed a significant correlation between mito-asymmetry and both R- and T-asymmetries of the nDNA (Fig. 6a). Our findings show that mutations prevalent in the mtDNA heavy strand are also common in the lagging and non-transcribed strands of the nDNA. Further analysis revealed that the correlations (Fig. 6a) are primarily driven by 6 base-specific asymmetries (Fig. 6c), not by their context (Supplementary Fig. 7-8). This indicates, that for example C>T mutations, regardless of context, occur more frequently than G>A across all three areas: (i) mtDNA heavy strand, (ii) nDNA lagging strand, and (iii) nDNA non-transcribed strand. Categorising the six mutation pairs by the degree of asymmetry, from highest to lowest, with the first mutation in each pair occurring more frequently, we got the next ranking: T>C, C>T, C>A, T>A, C>G, T>G (Fig. 6c). Interestingly two the most asymmetrical substitutions, C>T and A>G, are also the most common in the integral mtDNA mutational spectrum (Fig. 2a).

**Figure 6.**
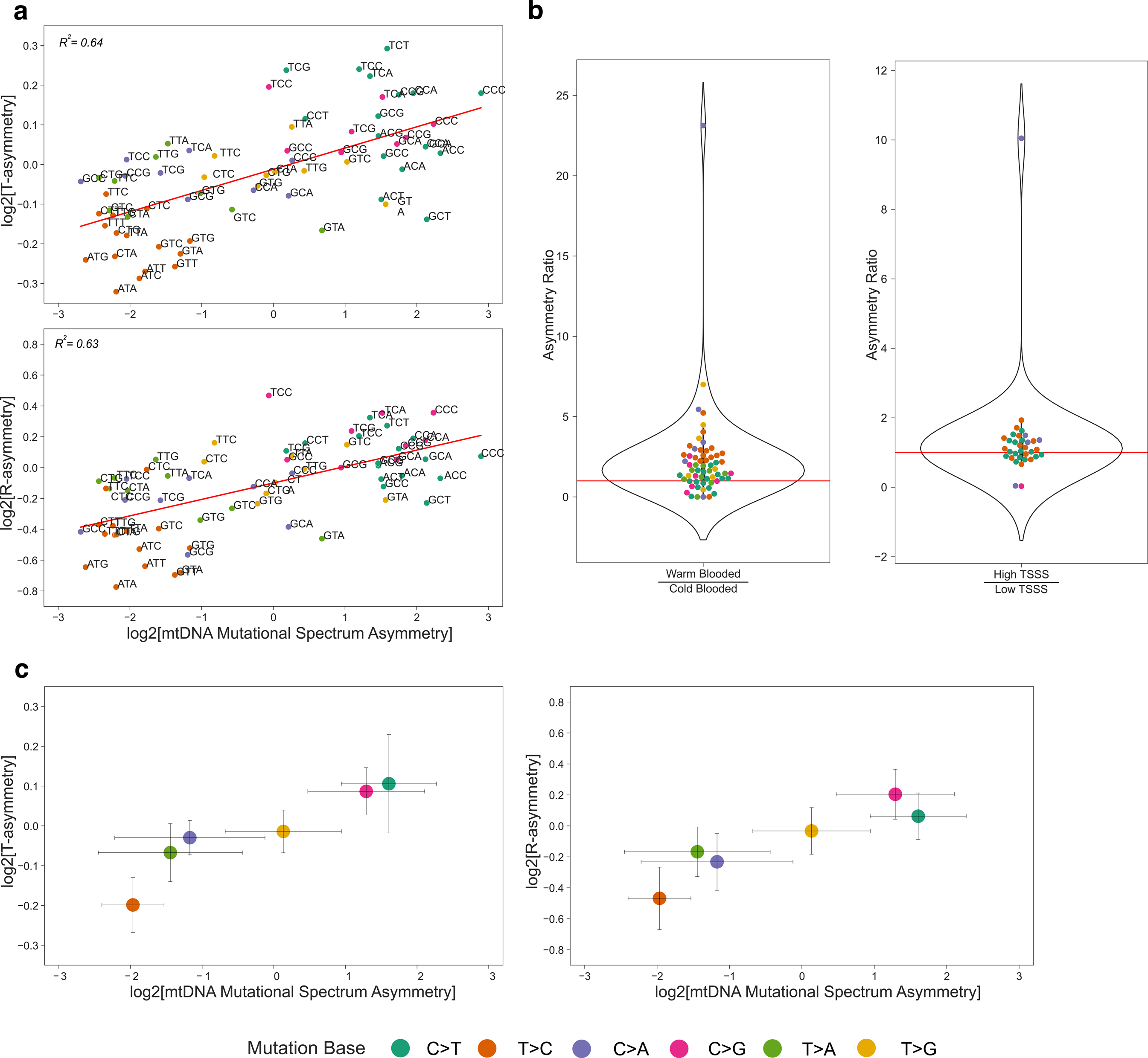
Comparison of the mitochondrial asymmetry based on the global mitochondrial mutational spectrum with the T and R asymmetries in nDNA. (a) The analysis of the mitochondrial asymmetry reveals a notable positive correlation with both T (upper panel, Spearman’s Rho =0.34, p = 0.0007, N = 96) and R (bottom panel, Spearman’s Rho = 0.29, p = 0.004, N = 96) nuclear asymmetries (each dot represents substitution type with a context). The elimination of zero-rated substitutions (rare transversions, never observed in chordates) from the mito-asymmetry significantly improved the positive associations with both T-(Spearman’s Rho = 0.64, p = 4.6*e-09, N = 68) and R-asymmetry (Spearman’s Rho = 0.62, p = 1.3*e-08, N = 68) (b) Direct comparison of mitochondrial asymmetries between warm- and cold-blooded species shows an increased strength of the asymmetry in warm-blooded (the ratio is higher than one) (Wilcoxon test, p = 1.56e-06, N=61). Similar trend is observed for high versus low TSSS regions (Wilcoxon test, p = 0.06, N=34). (с) Among the six primary substitution types, a robust association is observed between mito-asymmetry and T (left panel) and R (right panel) asymmetries, superseding the influence of nucleotide content.

What is the plausible mechanism for the asymmetry’s origin? The shared characteristic of the three areas described above is their single-stranded nature, suggesting a uniform mutational process influenced by ssDNA damage. If single-stranded specific damage^9,50,51^ is a viable hypothesis, we anticipate asymmetry to grow with (i) increased time spent single-stranded (TSSS) and (ii) increased total damage level. Assuming that TSSS is linearly increasing during asynchronous replication of mtDNA along the major arc^9^, we analysed human cancer mtDNA data^8^. Dividing the major arc into low and high TSSS zones (Methods) we revealed that asymmetry significantly increases in high TSSS areas, with the high to low TSSS asymmetry ratio exceeding one (Fig. 6b, right panel). Testing our hypothesis of different sources of mDNA damage, we leverage the convention that mitochondrial damage is linked to aerobic metabolic rates^7,52^: thus, mito-asymmetry in warm-blooded species should surpass that in cold-blooded ones. Comparing mito-asymmetry within the same gene (CytB) between the warmest (birds) and the coldest^34^ (fishes) chordata species in our dataset, we indeed observed an asymmetry ratio of warm to cold species greater than one (Fig. 6b, left panel).

While an alternative explanation for asymmetry, proposed for nDNA, involves low-fidelity translesion DNA synthesis (TLS) polymerases capable of error-prone bypassing of DNA lesions^4^, we find it less likely for mito-asymmetry. First, PrimPol, an error-prone TLS polymerase observed in mammalian mitochondria^53^, is rarely recruited and has a strong preference for generating base insertions and deletions^54^, which are rarely observed in mtDNA^8^. Second, the deamination of C and A on the heavy chain of mtDNA, resulting in the most common and asymmetrical mutations, C>T and A>G, are not expected to be helix-distorting changes that stall replication forks and necessitate PrimPol recruitment (see Zheng et al. 2006 ^55^). In fact, C>T substitutions are the most common among POLG-mediated errors in *in vitro* experiment^55^, suggesting that POLG can make C>T transitions via cytosine deamination without issues (Supplementary Fig. 9). In summary, we suggest that ssDNA damage is a key factor contributing to mutational asymmetry in mtDNA and potentially has some influence on nDNA as well.

## DISCUSSION

In this study, by integrating species-specific mtDNA mutational spectra from various chordates, we have reconstructed a comprehensive 192-component mutational spectrum. Our analyses deconvolute this spectrum into three main fundamental sources of mitochondrial mutations: replication-driven mutations and two damage-driven categories of mutations characterised by distinct etiologies and dynamics.

The first component, driven by POLG replication errors, comprises about 50% of all *de novo* mutations and includes C>T symmetrical mutations (SBS44 and SBS21 signatures, indicative of MMR deficiency in nDNA), A>G symmetrical mutations, and a symmetrical part of the majority of transversions (SBS5-like signature), shown in Fig. 7 in grey colour. The most prevalent type within this component is the symmetrical C>T substitutions, which also shows the most conservative pattern across chordate classes (see G_H_>A_H_ in Fig. 4c) and is known as the most common mutation for all polymerases including POLG due to the mispairing of T opposite to G, leading to C>T mutations^40,48^. If the frequency of symmetrical C>T (G_H_>A_H_) mutations and most transversions increases with each mtDNA replication round, we expect a positive correlation among mutation types in the component. Interestingly, this correlation has been confirmed recently by a principal component analysis in our comparative-species study, which shows collinearity between the majority of transversions and G_H_>A_H_ substitutions (Fig. 2c in Mihailova et al. 2022^10^).

**Figure 7.**
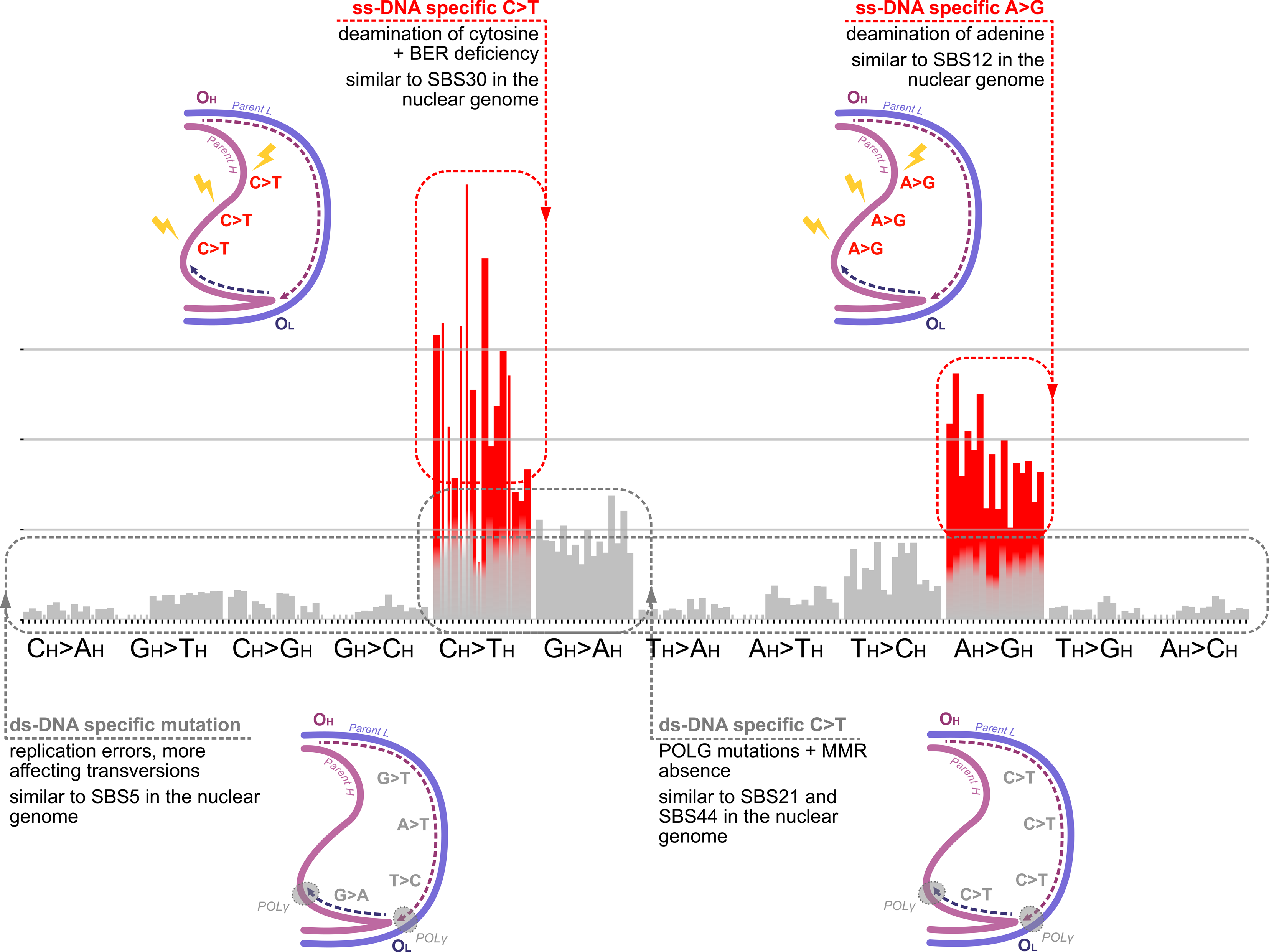
Graphical visualisation of the main mutational signatures in mtDNA: (i) symmetrical mutations, predominantly C>T, A>G and rare transversions (grey component), linked to POLG’s replication errors; similar to SBS21, SBS44 and SBS5; (ii) asymmetrical C>T mutations (red component), indicative of single-stranded DNA damage; similar to SBS30; (iii) asymmetrical A>G mutations (red component), also resulting from single-stranded DNA damage but particularly influenced by metabolic and age-specific mitochondrial environment; similar to SBS12.

The second component, representing roughly 30% of mutations and depicted in red in Fig. 7, comprises asymmetrical C>T mutations due to damage (SBS30 signature, indicating BER deficiency in nDNA). It is likely shaped by ssDNA damage caused by spontaneous deamination and oxidative stress, aggravated by deficient BER on the single-stranded heavy strand (Fig. 5a,c). The association of these mutations with dysfunctional NTHL1 in nDNA (Fig. 5), a glycosylase addressing oxidised pyrimidine lesions, and the prominent cCc>cTc signature (Supplementary Fig. 5a,c), experimentally confirmed as indicative of oxidative damage on ssDNA, suggest a significant oxidised component of these C>T mutations. Despite their high prevalence in mtDNA, these mutations maintain a rather constant rate across species, showing no sensitivity to metabolic or life-history changes^10,34^. This component’s insensitivity to metabolic or life-history traits and replication, highlighted by its distinct position relative to both the first component (G_H_>A_H_ mutations and the majority of transversions) and the third component (A_H_>G_H_) in our comparative species PCA plot (Fig. 2c in Mihailova et al. 2022^10^), suggests its potential as a molecular clock in mtDNA, similar to SBS1 in nDNA, which is characterised by C>T mutations in a CpG context due to deamination of 5-methylcytosine (SBS1). Although the methylation of cytosine in mtDNA remains uncertain and probably low^33^, somatic mtDNA mutations C>T in cancers clearly show a CpG context, indicating that formation of 5-methylcytosine and subsequent deamination may occur (Supplementary Fig. 5c, see Yuan et al. 2020^8^).

The third component, representing around 20% of the mutation spectrum, consists of asymmetrical A_H_>G_H_ mutations (depicted in red in Fig. 7), associated with ssDNA damage, primarily attributed to adenosine deamination. This mutation type, in contrast to the second component, exhibits notable correlations with eco-physiological traits in mammals^10^ and across chordates^49^. Its distinctiveness may be linked to its status as the most asymmetrical mutation in mtDNA, indicative of ssDNA damage. Moreover, chemical studies show that adenine is the most susceptible to deamination nucleotide^56^, and the process of deamination quickens at higher temperatures^57^ and in alkaline conditions^56^. This temperature dependency could account for the increased A_H_>G_H_ mutations and A_H_>G_H_ asymmetry, estimated as A_H_>G_H_/T_H_>C_H_ ratio, in warm-blooded compared to cold-blooded chordates^49^, while pH dependency might explain an association of A_H_>G_H_ with longevity and ageing in mammals^10^. All these findings, alongside the similarity of the third component to SBS12 and the potential implications of adenosine methylation, highlight important avenues for further research.

The categorisation of the mtDNA mutational spectrum into replication-driven, ssDNA damage-driven molecular clock-like and ssDNA damage-driven metabolism-associated components can enrich our understanding of species’ molecular evolution. It may reveal new aspects of species’ origins, history, and potential genetic factors influencing mutation rates, as it was shown in comparative studies on nDNA^2,58–61^. Additionally, these components offer insights into the dynamics of somatic mtDNA mutation patterns in different cancer and healthy tissues^62^.

Understanding the primary mutagens in mtDNA mutagenesis can guide strategies to reduce mtDNA mutation load, causative for many human diseases and ageing. For instance, evidence that damage contributes to up to 50% of de novo mutations suggests approaches to mitigate damage-induced mutations, either by reducing damage (for example by inducing hypoxia^29^) or by enhancing repair processes, like boosting mitochondrial BER via upregulation of NTHL1^44^. Further research is required to elucidate the damage processes impacting single-stranded mtDNA.

## Acknowledgements

This work was supported by the Federal Academic Leadership Program Priority 2030 at the Immanuel Kant Baltic Federal University (to D.I. and K.P.). A.G.M. and B.E. is supported by the Russian Science Foundation grant No. 21-75-20143. K.G. is supported by the Russian Science Foundation grant No. 21-75-20145. I.M. is supported by the Russian Science Foundation grant No. 21-75-10081. V.S. is supported by the Ministry of Science and Higher Education of the Russian Federation (agreement no. 075-15-2021-1084)

We thank the high-performance computing platform at the Immanuel Kant Baltic Federal University.

## Contributions

The design of the study developed by K.P. Data mining and processing performed by D.I., B.E. and K.G. Manuscript prepared by K.P., D.I., B.E. and A.G.M. All authors (D.I., B.E., A.G.M., V.S., M.K.S., I.M., D.K., W.S.K., P.K., S.D., K.K., J.F., K.G. and K.P.) discussed in depth the manuscript and the rationale behind the project. All authors read and approved the final manuscript. The authors express their appreciation to Vladimir Seplyarskiy for his valuable contributions and insightful discussions regarding asymmetry and repair mechanisms.

