## Supplementary Materials for "Mitochondrial mutation spectrum in Chordates: damage versus replication signatures, causes, and dynamics"

#### 1. Variations in mtDNA Mutational Spectrum within Chordates: Importance of Damage

**Supplementary Table 1. The table of calculated mutational spectra for all available 967 chordate species.**

https://github.com/mitoclub/mtdna-192component-mutspec-chordata/blob/main/ToPaper/SuplFiles/SupplementaryTable1.csv

*Example:*

| Species | Class | A[A>G]A | T[T>C]T | T[T>G]C |
|---|---|---|---|---|
| Abbottina Rivularis | Actinopterygii | 0.002474 | 0.005 | 0.000 |
| Ambystoma Barbouri | Amphibia | 0.009 | 0.003 | 0.000 |
| Mus Musculus | Mammalia | 0.012 | 0.002 | 0.002 |

*Species - Name of vertebrate species in latin for which the 192-component mitochondrial mutation spectrum was calculated*
*Class - Class of vertebrate animal out of five possible: actinopterygii, amphibia, lepidosauria, mammalia, aves*
*Next columns present the unique substitutions with the normalized substitution rate for them*
*All the values of the substitution rates are rounded to 3 decimal places.*

**Supplementary Table 2. Calculated 192-component mutational spectrum of vertebrate classes, vertebrates together, and human cancers in COSMIC format.**

https://github.com/mitoclub/mtdna-192component-mutspec-chordata/blob/main/ToPaper/SuplFiles/SupplementaryTable2.csv

We provide a comprehensive overview of the normalized substitution rates for different types of substitutions in five classes of vertebrates, as well as the combined analysis for all vertebrates and human cancers. All the values of the substitution rates are rounded to 4 decimal places

*Example:*

| Type | Actinopterygii | Amphibia | Lepidosauria | Mammalia | Aves | Cancers | Vertebrate |
|---|---|---|---|---|---|---|---|
| A[A>G]A | 0.0107 | 0.0101 | 0.0111 | 0.0167 | 0.0423 | 0.0083 | 0.0145 |
| T[A>C]A | 0.0005 | 0.0002 | 0.0004 | 0.0002 | 0.0009 | 0.0000 | 0.0004 |
| G[A>C]A | 0.0006 | 0.0003 | 0.0006 | 0.0008 | 0.0006 | 0.0000 | 0.0006 |

**Supplementary Fig. 1. Estimation of synonymous sites neutrality.**

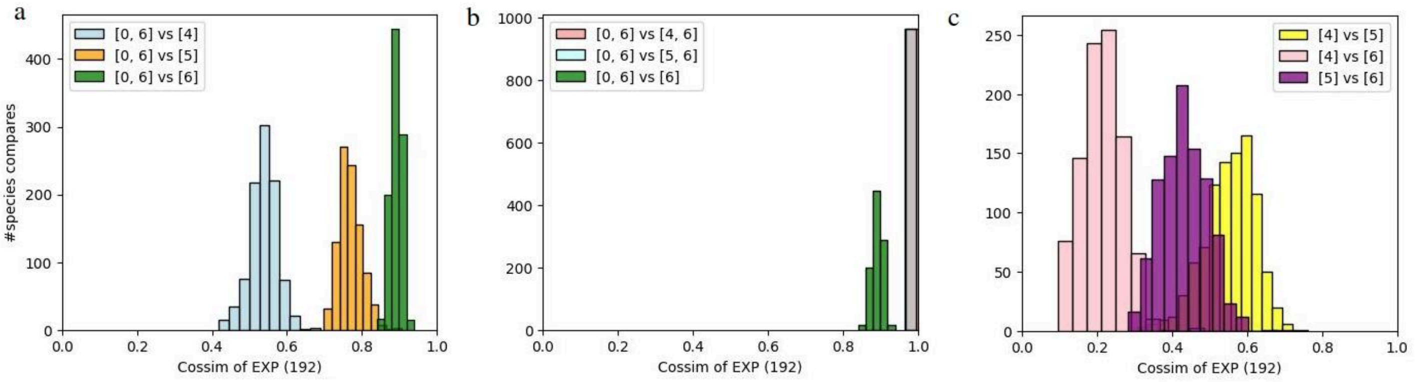


To assess the neutrality of synonymous sites in mtDNA, we compared expected substitutions counts (192-component vectors) in different sites samples according to estimated site variability rate for each species. The legend delineates the comparison between site samples based on their rate categories. For instance, "[0, 6] vs [5]" indicates a comparison of the expected substitution counts across all sites encompassing the full range of rate categories from 0 to 6, against the substitution counts from sites categorized exclusively under rate category 5. To estimate rate heterogeneity among sites, we utilized the IQ-TREE v2.2.0[1] software for general phylogenetic inference with the specified options (-m 10.12+FO+G6+I -rate) on codon alignment of reference CytB sequences of vertebrates species. The options means that all sites were divided into 6 rate categories, reflecting a gradient of variability from low to high, plus zero-category for invariable sites.

Our analysis selectively focused on sites with rate categories 4, 5, and 6, since possible (expected) synonymous substitutions in other categories were represented by a fraction of less than 1%. Figures a and b show that expected substitution counts on sites with different rates ([4, 6], [5, 6], [6]) highly similar to substitution counts on all sites [0, 6], i.e. exclusion of conservative sites did not significantly alter the expected substitutions counts as well as the mutational spectra. However, future analyses may unveil the nature of highly-constrained sites, whether they result from selection and should be excluded in reconstructing the neutral mutational spectrum or if they represent mutational cold spots, which can be tested in longer nucleotide contexts.

**Supplementary Fig. 2. Pairwise cosine similarities between all species. Colorbar indicates cosine similarity.** Species were hierarchically clustered inside each class, accordingly, they were ordered by cosine similarity in each class. Despite the red areas on the heatmap that indicate small clusters, it is obvious that species from different classes are as similar as species within the class, i.e there is no clusterization of species spectra into classes.

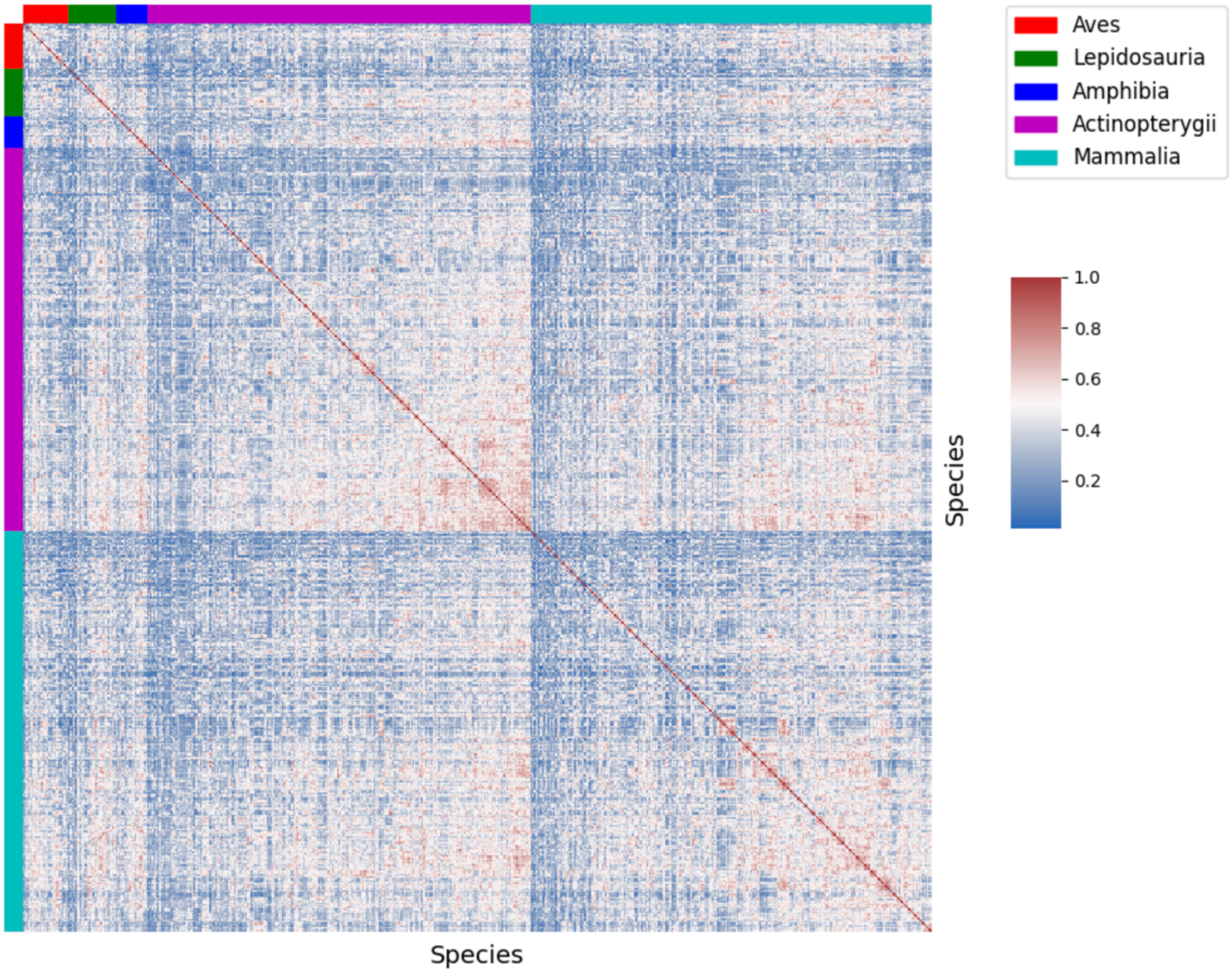

### Supplementary Fig. 3. Separate class-specific spectra for each of the five chordata classes and cancer data

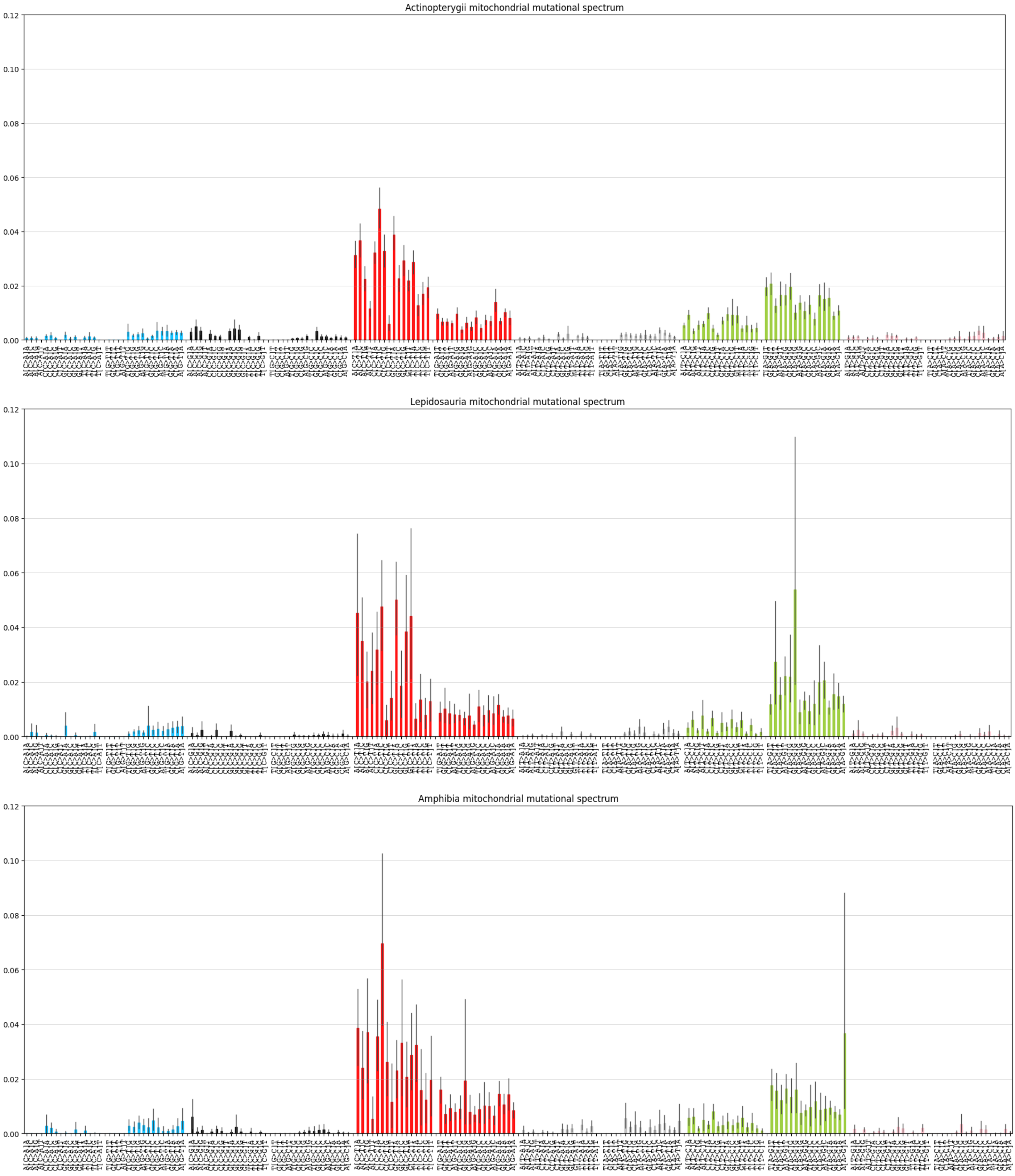

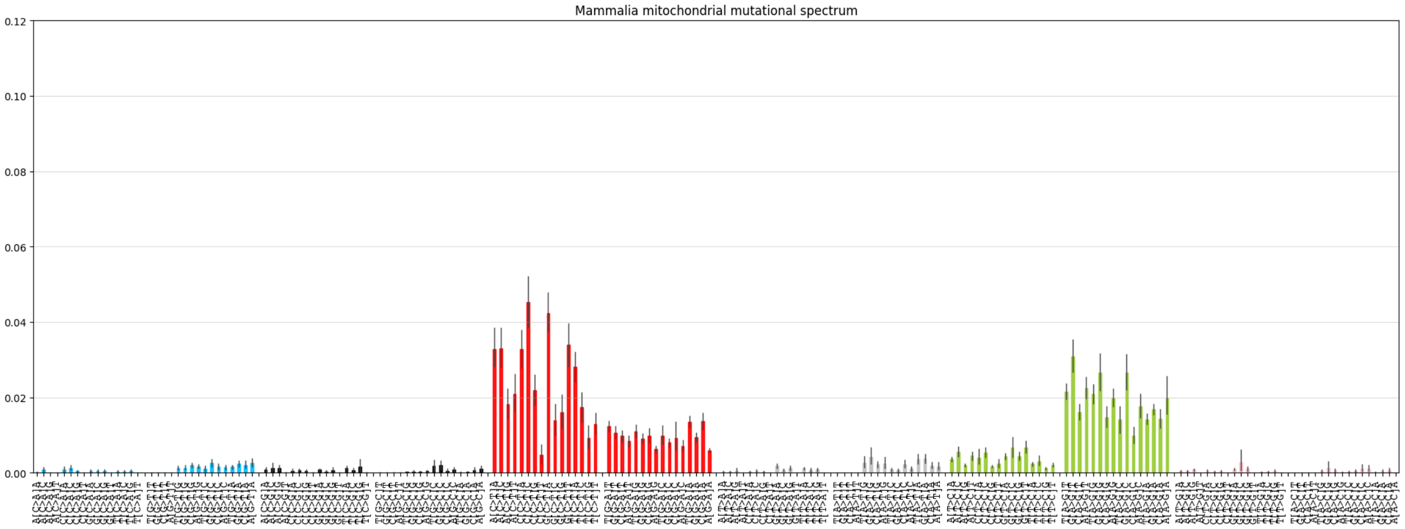
Mammalia mitochondrial mutational spectrum

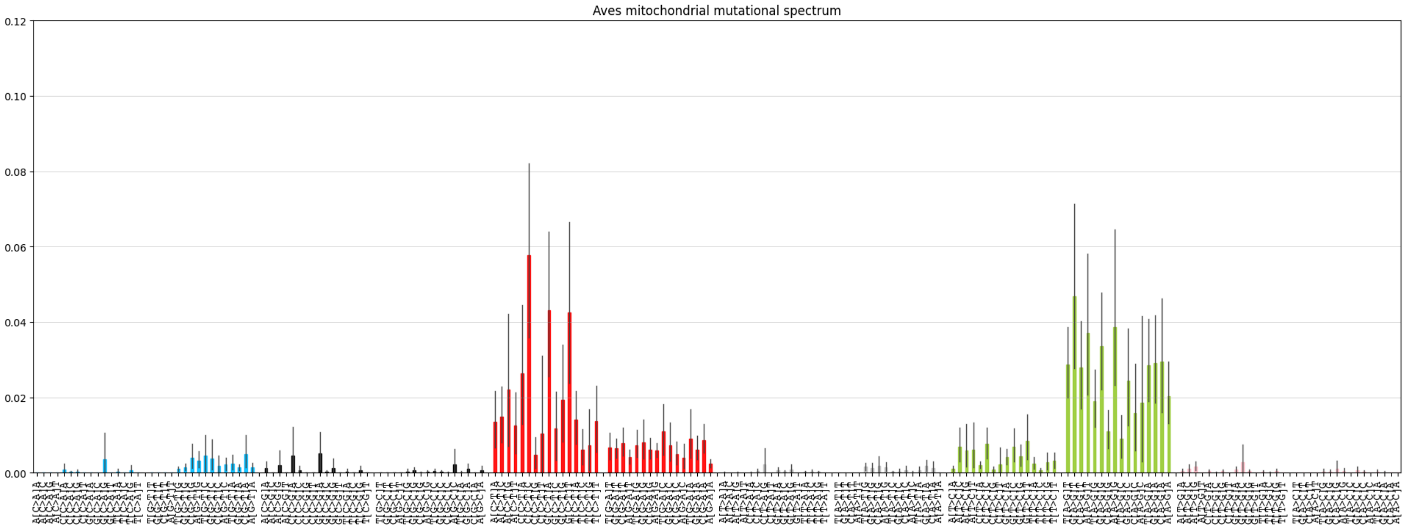
Aves mitochondrial mutational spectrum

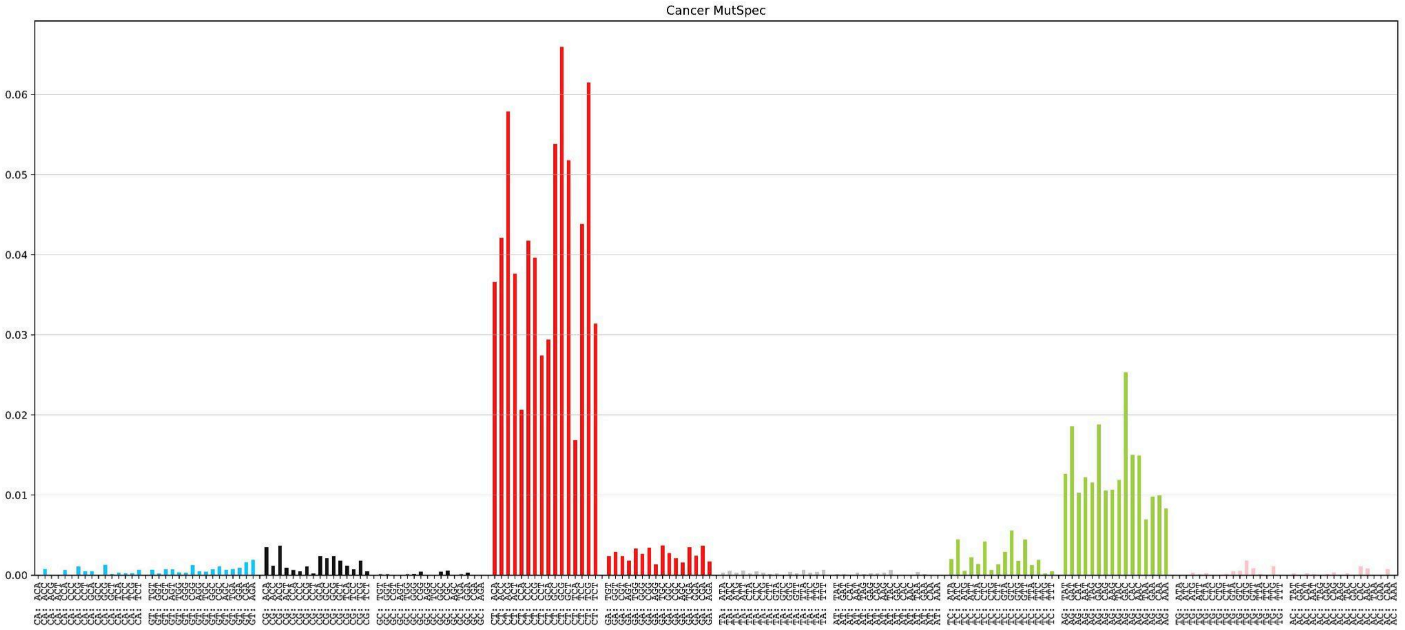
Cancer MutSpec

#### 2. Conservative and variable patterns of the mtDNA spectrum: symmetrical replication-driven and asymmetrical damage-driven mutations

**Supplementary Fig. 4. Lack of transversions compared to transitions in chordate spectra**

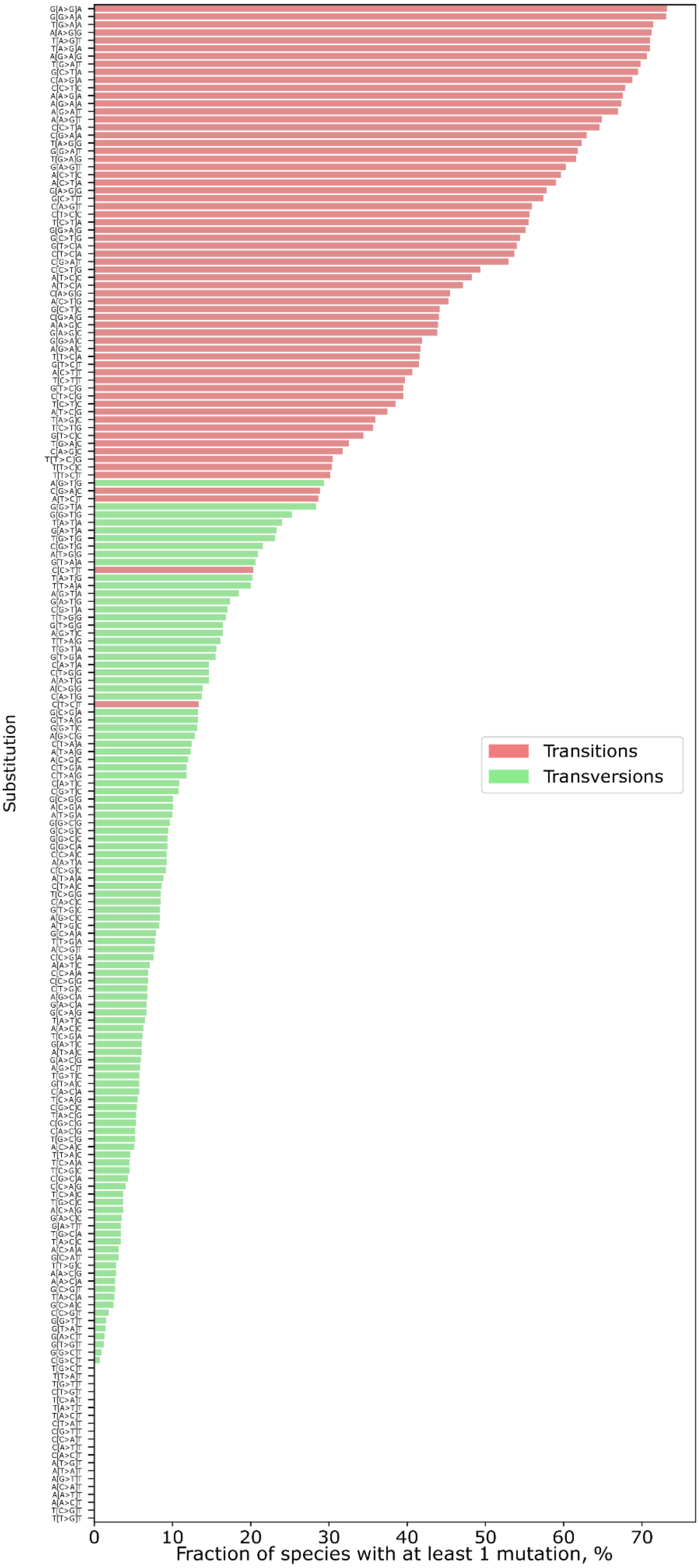

**Supplementary Fig. 5. $C_H$>$T_H$ and $A_H$>$G_H$ motifs in mtDNA mutational spectrum in chordates and human cancers**

*Full version with chordata classes:*

*For C>T context:*

*https://github.com/mitoclub/mtdna-192component-mutspec-chordata/blob/main/ToPaper/SupplementaryFigure5_CTcontext.pdf*

*For A>G context:*

*https://github.com/mitoclub/mtdna-192component-mutspec-chordata/blob/main/ToPaper/SupplementaryFigure5_AGcontext.pdf*

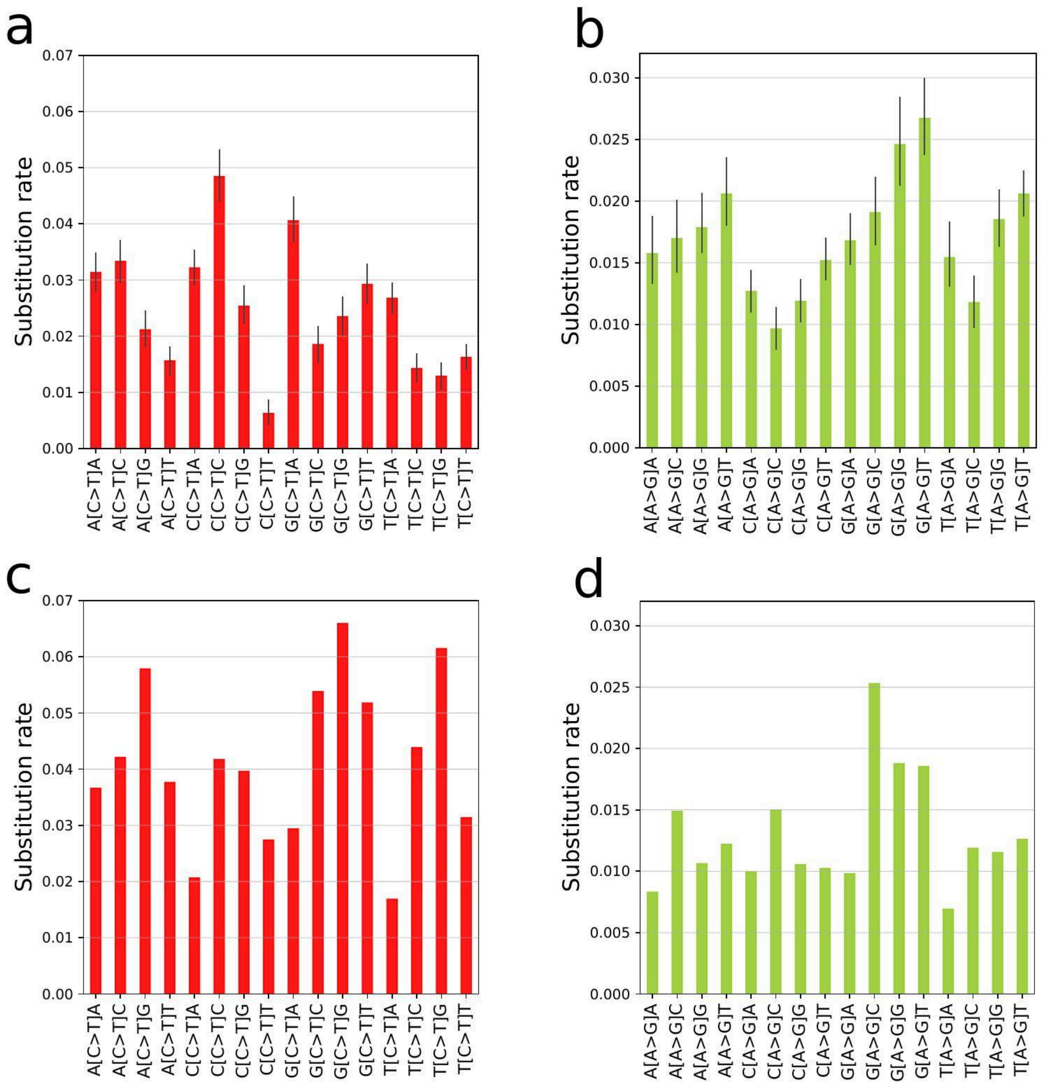


(a) $C_H$>$T_H$ motifs in chordate mtDNA spectrum; the motif cCc>cTc is consistently the most frequently observed motif. This pattern was previously reported as the hallmark of oxidative damage in single-stranded DNA. (b) $A_H$>$G_H$ motifs in chordate mtDNA spectrum; nAt>nGt and nAg>nGg are the second two of the most common motifs. It has been reported that such motifs can be linked to the 6mA ROS induced mutations in mtDNA. (c) $C_H$>$T_H$ spectrum motifs presented in cancer data; asymmetrical $C_H$>$T_H$ substitutions are most frequent in the context of nCg. This suggests two potential explanations: at first, there can be reduced oxidative damage affecting somatic cancer mtDNA mutations compared to germ-line mtDNA mutations, leading to a decrease in cCc>cTc. Secondly, the potentially stronger role of cytosine methylation in somatic mtDNA mutagenesis can result in an increase in CpG > TpG. While being less explored in the context of mtDNA, this phenomenon may be more prominent in somatic tissues echoing the well-known association of CpG > TpG substitutions with ageing and molecular evolution in the nuclear genome. (d) $A_H$>$G_H$ spectrum motifs presented in cancer data.

3. ***mtDNA mutations through the lens of COSMIC signatures: BER deficiency and*** $C_H$>$T_H$, ***MMR absence and*** $G_L$>$A_L$, ***and SBS12-driven*** $A_H$>$G_H$ ***alterations***

**Supplementary Fig. 6. Splitting of the asymmetrical mitochondrial 192-component spectrum into three sets for signatures deconvolution analysis**

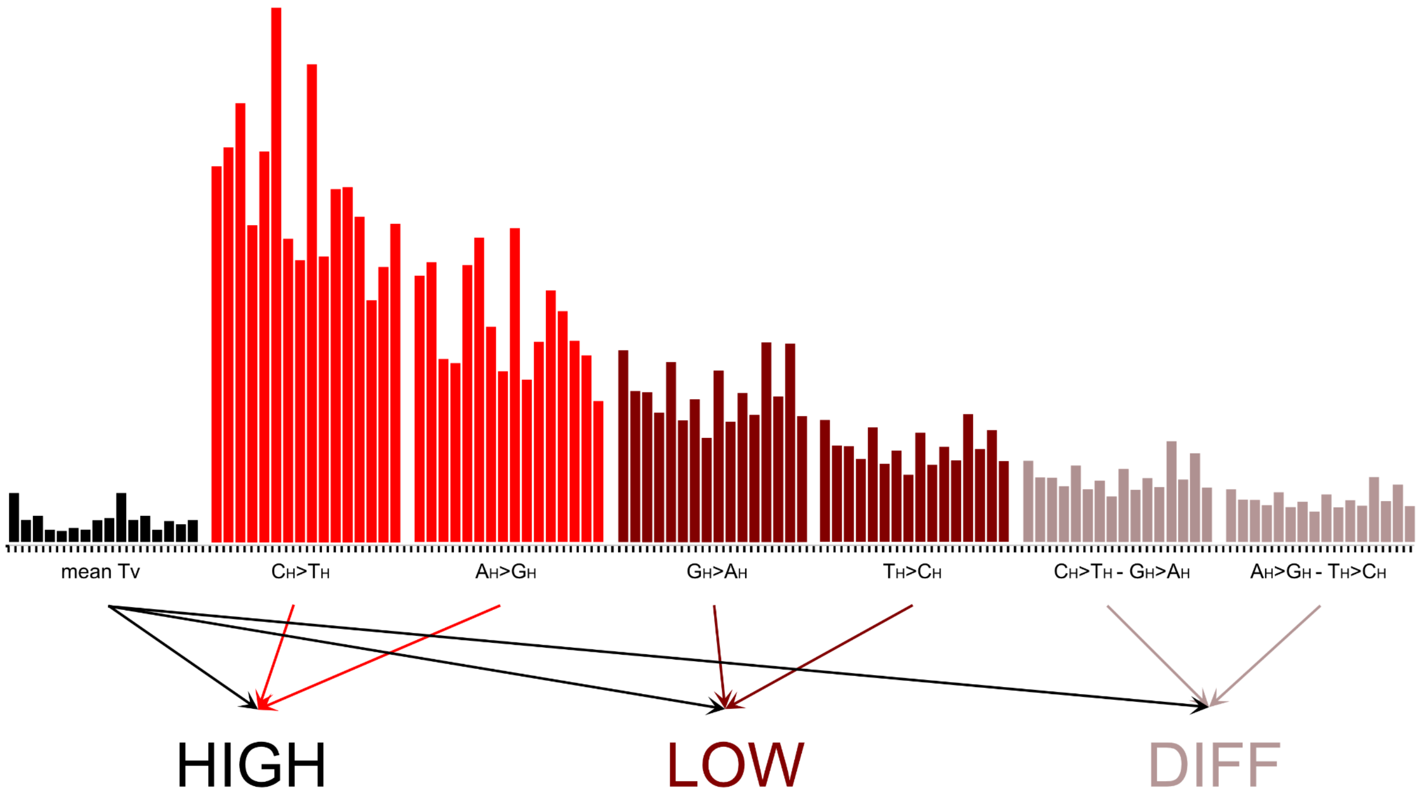


The "high" spectra include more frequent $C_H$>$T_H$ and $A_H$>$G_H$ transitions on heavy strand, while the "low" spectra include $G_H$>$A_H$ and $T_H$>$C_H$ transitions on heavy strand; “diff” spectra calculated by subtracting “low” from “high” spectra in a context-dependent manner. All complementary pairs of transversions rates were averaged (mean Tv) and equally added to the "low", “diff” and "high" spectra.

4. **Strong asymmetry in mtDNA mutagenesis is shaped by single-strand DNA damage coupled with Base Excision Repair deficiency during asynchronous replication**

**Supplementary Fig. 7. Association of the nuclear T asymmetry and asymmetry of the six main types of substitutions in mitochondrial mutational spectrum for all chordates**

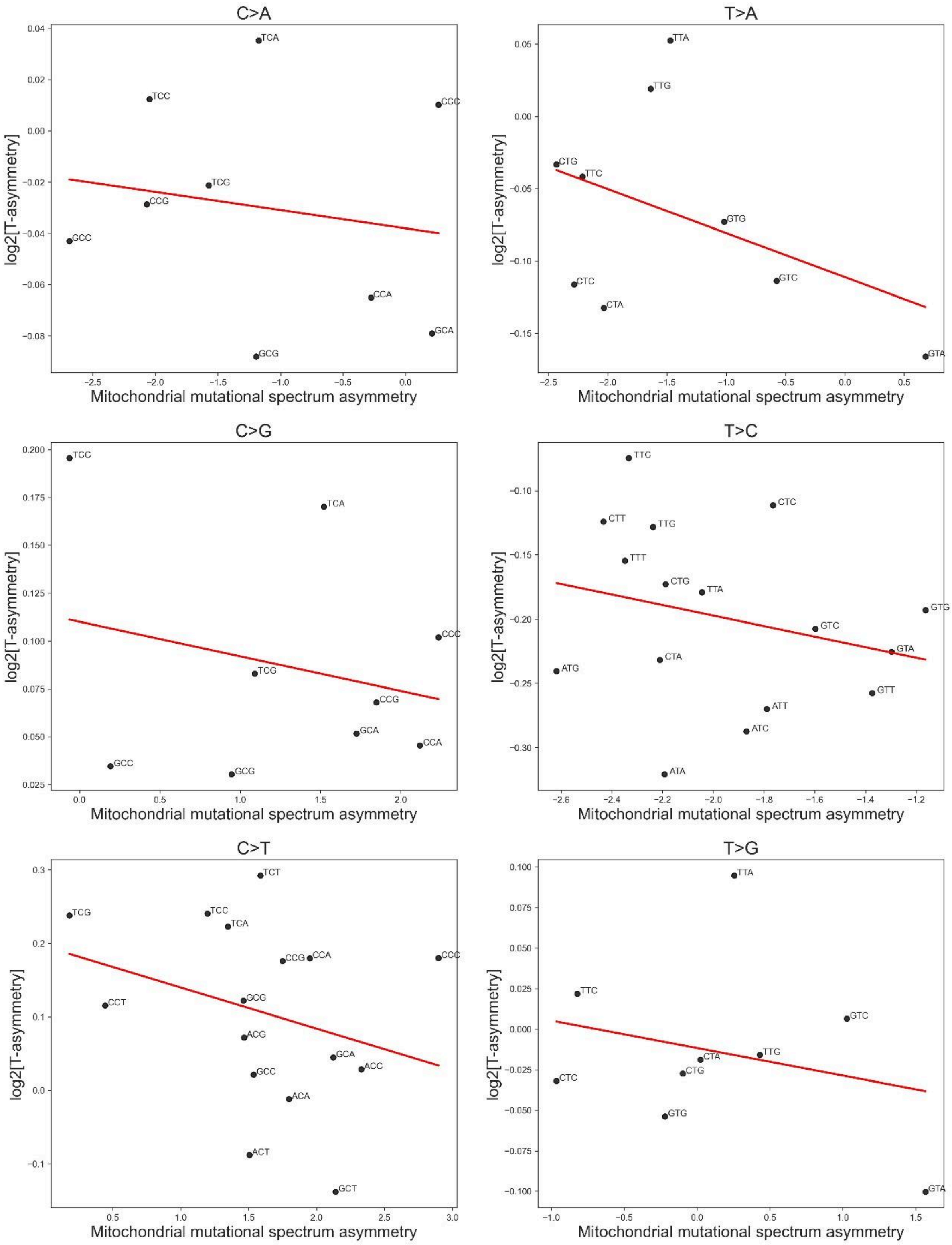

**Supplementary Fig. 8. Association of the nuclear R asymmetry and asymmetry of the six main types of substitutions in mitochondrial mutational spectrum for all chordates**

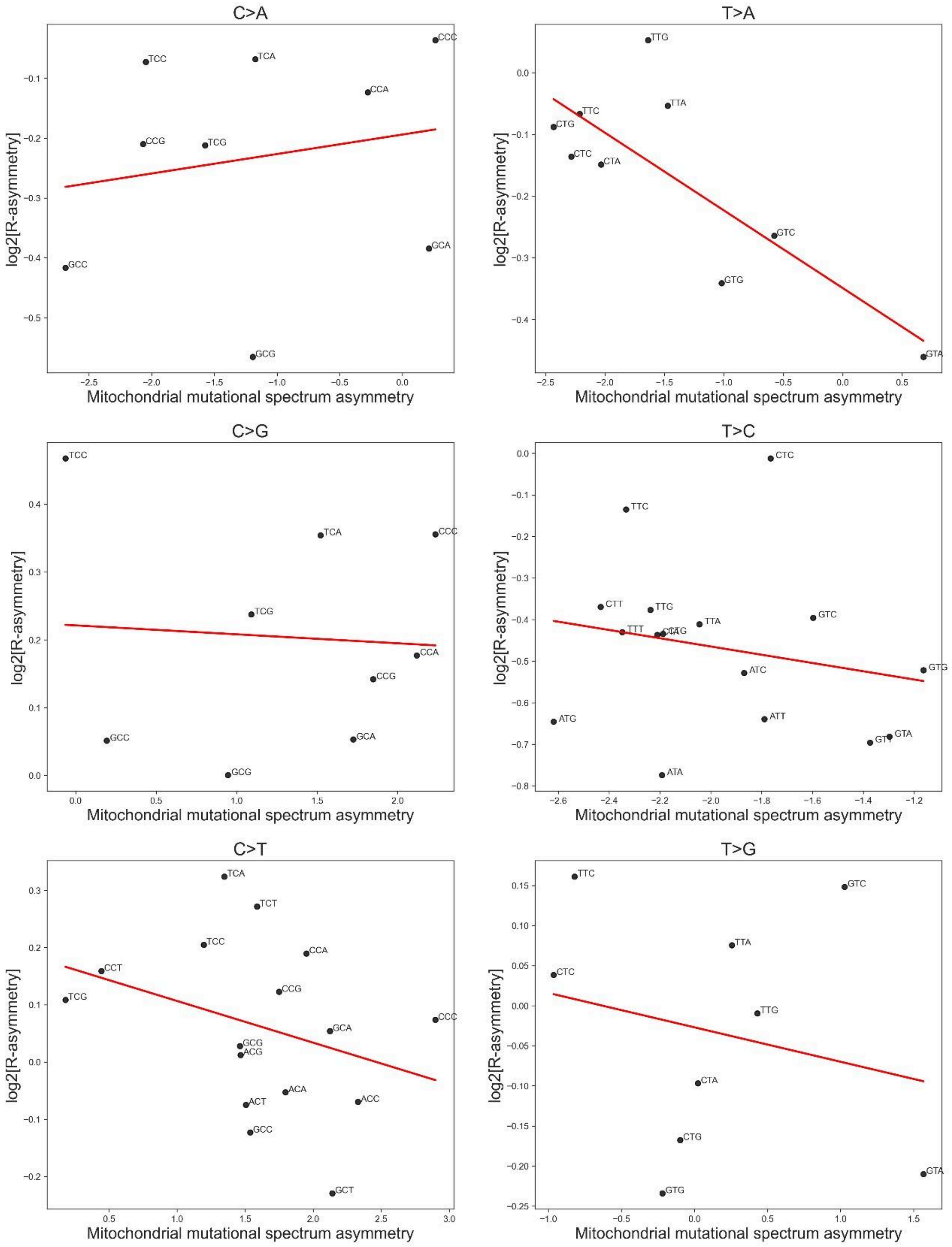

**Supplementary Fig. 9. Symmetrical mutagenesis of mtDNA shaped by POLG**
*https://github.com/mitoclub/mtdna-192component-mutspec-chordata/blob/main/scripts/PolGspectrum.Rmd*

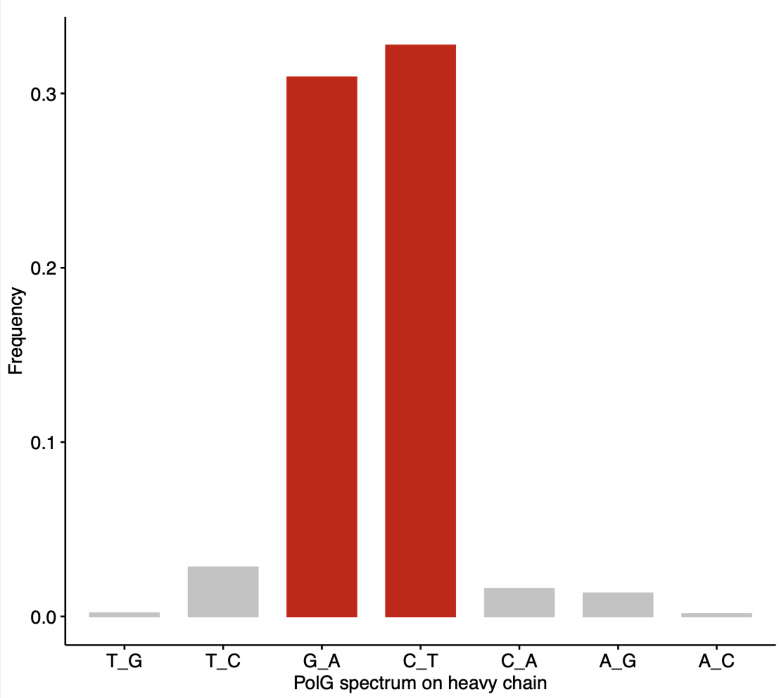


Recalculation of POLG' mutational spectrum based on experimental data [2]. We demonstrate that normalized rates of $C_H$>$T_H$ and $G_H$>$A_H$ mutations are rather the same, confirming the symmetrical nature of mutations, mediated by POLG via cytosine deamination.
